# Development of Recombinant Anti-TLR2 Antibodies and PLGA Nanoparticle-based Gene Therapy for the Treatment of Neuropathic Pain

**DOI:** 10.1101/2025.11.13.688281

**Authors:** Subeen Lee, Jaekyung Jeon, Hyunji Lee, Ellane Eda Barcelona, Jinpyo Hong, Sung Joong Lee

## Abstract

Neuroinflammation is a key contributor to neuropathic pain, with microglial Toll-like receptor 2 (TLR2) playing a central role in initiating and sustaining proinflammatory responses. However, existing TLR2-targeting antibody therapies are limited by poor delivery to the central nervous system and short-lived efficacy. We developed a non-viral gene therapy strategy using biodegradable poly(lactic-co-glycolic acid) (PLGA) nanoparticles (NPs) to deliver recombinant anti-TLR2 antibody genes. High-affinity nanobody and single-chain variable fragment candidates were selected through phage display screening and shown to suppress TLR2-dependent signaling both *in vitro* and *in vivo*. In mouse models of neuropathic pain, a single intrathecal administration of PLGA NP–encapsulated antibody genes produced robust and sustained analgesia, accompanied by reduced glial activation and proinflammatory cytokine expression. These findings demonstrate a modular NP-based platform for sustained antibody expression in the central nervous system and establish its potential for the treatment of chronic pain driven by innate immune activation.

## INTRODUCTION

Neuropathic pain is a chronic pain condition characterized by spontaneous pain, allodynia (pain hypersensitivity to normally innocuous stimuli), and hyperalgesia (amplified pain response to noxious stimuli), often resulting from peripheral nerve injury.^1^ Neuropathic pain has an estimated global prevalence of 7–10% in the general population and up to 30% in diabetic patients.^2^ Despite that high prevalence and societal burden, effective long-term pharmacological treatments remain limited. Current therapeutic approaches for neuropathic pain primarily comprise anticonvulsants (e.g., gabapentin, pregabalin), antidepressants (e.g., duloxetine, amitriptyline), and opioids (e.g., tramadol, tapentadol).^3^ These drugs primarily act on neuronal excitability and neurotransmitter signaling to alleviate symptoms, but they do not address the underlying pathophysiology. Furthermore, their clinical use is often restricted by adverse effects such as sedation, dizziness, weight gain, and (in the case of opioids) tolerance and dependence. Such effects are mainly due to off-target toxicity, lack of specificity, and the need for repeated dosing because of their short half-lives.^4^ Thus, there is an urgent need for novel therapeutic strategies that target the core mechanisms driving disease progression and offer improved safety, durability, and accessibility.

Chronic or excessive neuroinflammation underlies various central nervous system (CNS) disorders, including neuropathic pain.^5^ Neuroinflammation is a physiological defense system. Following an insult, activated microglia and astrocytes release proinflammatory cytokines (e.g., TNF-α, IL-1β, IL-6), chemokines (e.g., CCL2), and reactive molecules such as nitric oxide and reactive oxygen species.^6^ Those mediators promote pathogen clearance, recruit peripheral immune cells, and facilitate debris removal and tissue repair. However, in pathological conditions, glial cells remain activated, leading to sustained secretion of proinflammatory mediators and oxidative stress, which contribute to central sensitization and hypersensitivity.

Microglia play a central role in initiating and sustaining neuroinflammation, largely through pattern recognition receptors such as Toll-like receptors (TLRs).^7,8^ Among them, TLR2 is highly expressed on microglia and recognizes pathogen-associated molecular patterns (PAMPs) and damage-associated molecular patterns (DAMPs), including bacterial lipoproteins, peptidoglycans, and endogenous molecules released during cellular stress or nerve injury.^9,10^ Neuropathic pain is a prototypical example in which microglial TLR2-mediated inflammatory responses play a critical role in disease progression. TLR2 is upregulated and activated in spinal microglia after nerve injury, acting as an initiator of the inflammatory cascade^11^. Notably, TLR2-knockout mice exhibit attenuated nociceptive behaviors following peripheral nerve injury, and the upregulation of NADPH oxidase-2 and other inflammatory genes is abolished.^11,12^ Those findings establish microglial TLR2 as a key mediator of neuroinflammation and a promising target for therapeutic intervention in neuropathic pain.

Therefore, we aimed to develop a novel therapeutic strategy targeting microglial TLR2 for the treatment of neuropathic pain. To overcome the limitations of current therapies, we produced recombinant antibodies—nanobodies (VHH) and single-chain variable fragments (scFvs), two widely used recombinant antibody formats—as alternatives to conventional full-length immunoglobulins (IgGs).^13,14^ Nanobodies are derived from the heavy-chain-only antibodies found in camelids and comprise a single monomeric variable domain. scFvs are engineered by genetically linking the variable heavy (VH) and variable light (VL) domains of an antibody via a short flexible peptide linker. These small antibodies (nanobody: ∼15 kDa; scFv: ∼25 kDa) exhibit excellent stability, target specificity, low immunogenicity, good genetic modularity, simple production, and compatibility with high-throughput screening.^13,15–17^ We successfully found TLR2-binding nanobodies and scFvs through phage display biopanning, and we have demonstrated that they exhibit greater analgesic effects in a mouse neuropathic pain model than conventional full-length IgG antibodies.

Furthermore, to enhance their effect and duration, we used a gene therapy–based approach. Traditional gene therapy approaches have predominantly relied on viral vectors, such as adeno-associated virus (AAV). Although those vectors are effective in achieving high transduction efficiency and prolonged gene expression, they have significant limitations, including the potential for insertional mutagenesis (particularly with integrating viruses) and dose-limiting immune responses that hinder re-administration.^18,19^ Recent reports of AAV-associated hepatotoxicity in patients with Duchenne muscular dystrophy underscore the safety concerns that continue to challenge the broad application of viral gene delivery systems.^20^

In response to the limitations of AAV, non-viral gene delivery platforms have gained increasing attention. Among the non-viral carriers, poly(lactic-co-glycolic acid) (PLGA) nanoparticles (NPs) have emerged as a delivery platform widely used across various therapeutic areas due to their excellent biocompatibility and biodegradability.^21^ In this study, we demonstrate that gene delivery via a single intrathecal injection of PLGA NPs enabled long-term expression of the therapeutic recombinant antibodies in mouse spinal cords. Notably, in a mouse model of neuropathic pain, this approach induced analgesia that lasted up to 8 weeks and attenuated glial reactivity. Together, our results suggest the therapeutic potential of combining recombinant antibody engineering with PLGA NP–based gene delivery to target microglial TLR2–driven neuroinflammation.

## RESULTS

### Screening and identification of anti-TLR2 recombinant antibody candidates

To screen for novel TLR2-neutralizing nanobodies and scFvs by phage display library screening, we first constructed a synthetic nanobody library using polymerase chain reaction (PCR)-based gene assembly (Figure S1A). The nanobody scaffold was based on hs2dAb, a humanized, synthetic, single-domain antibody from a previous study (Figure 1A).^22^ The complementarity-determining regions (CDRs) within the primers contained randomized nucleotide sequences to introduce antibody diversity. In parallel with the nanobody library, an scFv antibody library was also prepared to expand the antibody format diversity. The scFv library from Creative Biolabs was constructed using VH and VL fragments derived from human peripheral blood mononuclear cells.

**Figure 1.**
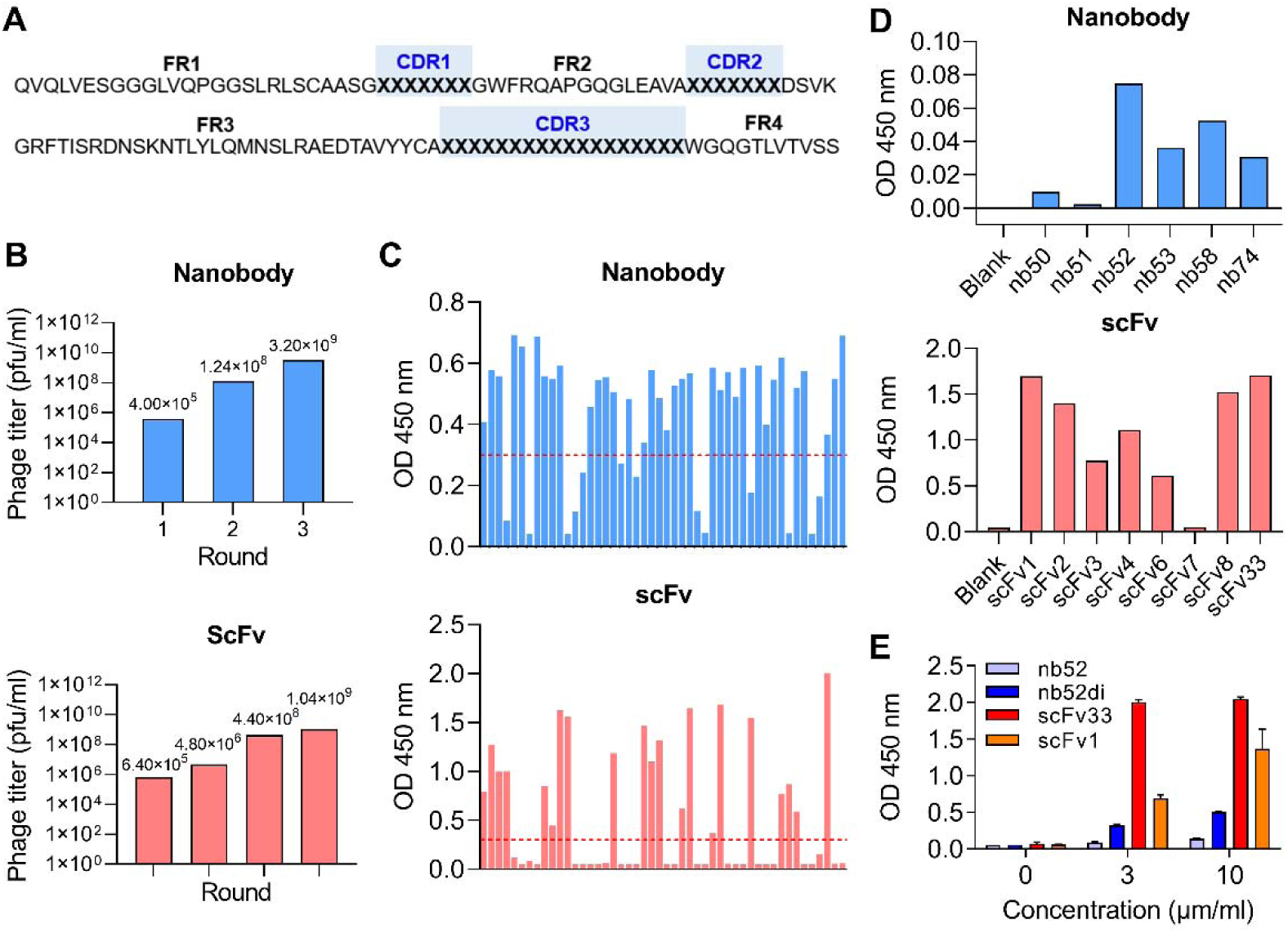
Screening of anti-TLR2 recombinant nanobodies and scFvs. **(A)** Amino acid sequence used for nanobody design, with randomized CDRs highlighted in blue. **(B)** Phage titers measured after each round of phage display biopanning in the nanobody and scFv libraries. **(C)** ELISA screening of the individual phage-displayed clones obtained and amplified after the final round of biopanning. The red dashed lines indicate the OD_450_ threshold of 0.3. See also Figure S1D. **(D)** ELISA analysis using crude antibody preparations from the nanobody and scFv candidate clones. **(E)** Concentration-dependent ELISA of four selected antibodies—nb52, its dimeric form (nb52di), scFv33, and scFv1—using crude antibody preparations.

To generate a phage display library for antibody screening, the recombinant antibody gene fragments were cloned into the pADL23c phagemid vector (Figure S1B). The phagemids were introduced into TG1 *Escherichia coli*, producing a diverse pool of phages that displayed recombinant antibodies fused to pIII on their surfaces. To isolate TLR2-specific antibodies, both the nanobody and scFv phage libraries were subjected to multiple rounds of phage display biopanning using a recombinant TLR2-coated plate (Figure S1C). Phage titers of the eluted fractions were measured at each round to monitor the enrichment of TLR2-specific binders. A progressive increase in phage titers was observed across the rounds, from ∼10 pfu/mL in the first round to ∼10 pfu/mL in the final round for both libraries (Figure 1B). This robust enrichment indicates our successful selection and amplification of high-affinity anti-TLR2 binders. After the final rounds of biopanning, the enriched phage pools were individually amplified and evaluated by ELISA for their TLR2-binding activity (Figures 1C and S1D). Clones with OD_450_ values ≥ 0.3, indicative of strong binding to TLR2, were selected for sequence analysis. The results revealed that certain antibody sequences appeared repeatedly within the selected clones (Table 1), indicating convergence toward highly specific TLR2-binding motifs.

Based on the ELISA data and the frequency of recurring sequences, crude antibody extracts of candidate TLR2-specific binders were prepared from induced *E. coli* cultures. The TLR2 reactivity of the extracts was then assessed by ELISA (Figure 1D). Among the nanobody candidates, nb52 exhibited the strongest signal (OD ≈ 0.08). To improve its TLR2-binding affinity, a divalent version of nb52 was generated by genetically linking two identical VHH domains in tandem. Among the scFv candidates, scFv33 and scFv1 showed markedly strong binding, with OD values exceeding 1.5. To further compare the binding performance of the final antibody candidates—monovalent and divalent forms of nb52 and scFv33 and scFv1—a concentration-dependent ELISA was performed using crude proteins (Figure 1E). All the antibodies showed a dose-dependent increase in signal. Notably, the nb52 dimer exhibited better binding than its monovalent form (OD at 10 μg/mL: ∼0.5 vs. ∼0.2, respectively). Among all the tested candidates, scFv33 showed the strongest binding to TLR2, with an OD value of ∼2.0 at 10 μg/mL.

In summary, the *in vitro* phage display workflow efficiently identified high-affinity anti-TLR2 antibody candidates. Four lead clones—nb52, nb52 dimer, scFv33, and scFv1—were selected based on their binding strength and sequence convergence. These results validate the recombinant antibody screening pipeline as a time– and cost-efficient strategy for antibody discovery.

### Biochemical and functional characterization of anti-TLR2 nanobodies and scFvs

We next assessed the biochemical and functional properties of the selected anti-TLR2 antibody clones. To this end, we obtained purified antibody proteins. SDS-PAGE and western blot analyses confirmed the successful expression and purification of the nb52 monomer, nb52 dimer, scFv33, and scFv1, with bands detected at the expected molecular weights (Figure S2).

Surface plasmon resonance (SPR) analyses revealed dose-dependent binding of all four antibodies to immobilized TLR2, with scFv33 showing the highest affinity (K_D_ = 111 nM), followed by scFv1 (131 nM), nb52 dimer (167 nM), and nb52 monomer (1510 nM) (Figure 2A). Notably, both the nb52 dimer and scFv33 showed slower dissociation rates as well as lower K_D_ values, indicating more stable binding to TLR2. No measurable binding was observed to TLR3 or TLR4 (Figure S3), confirming specificity.

**Figure 2.**
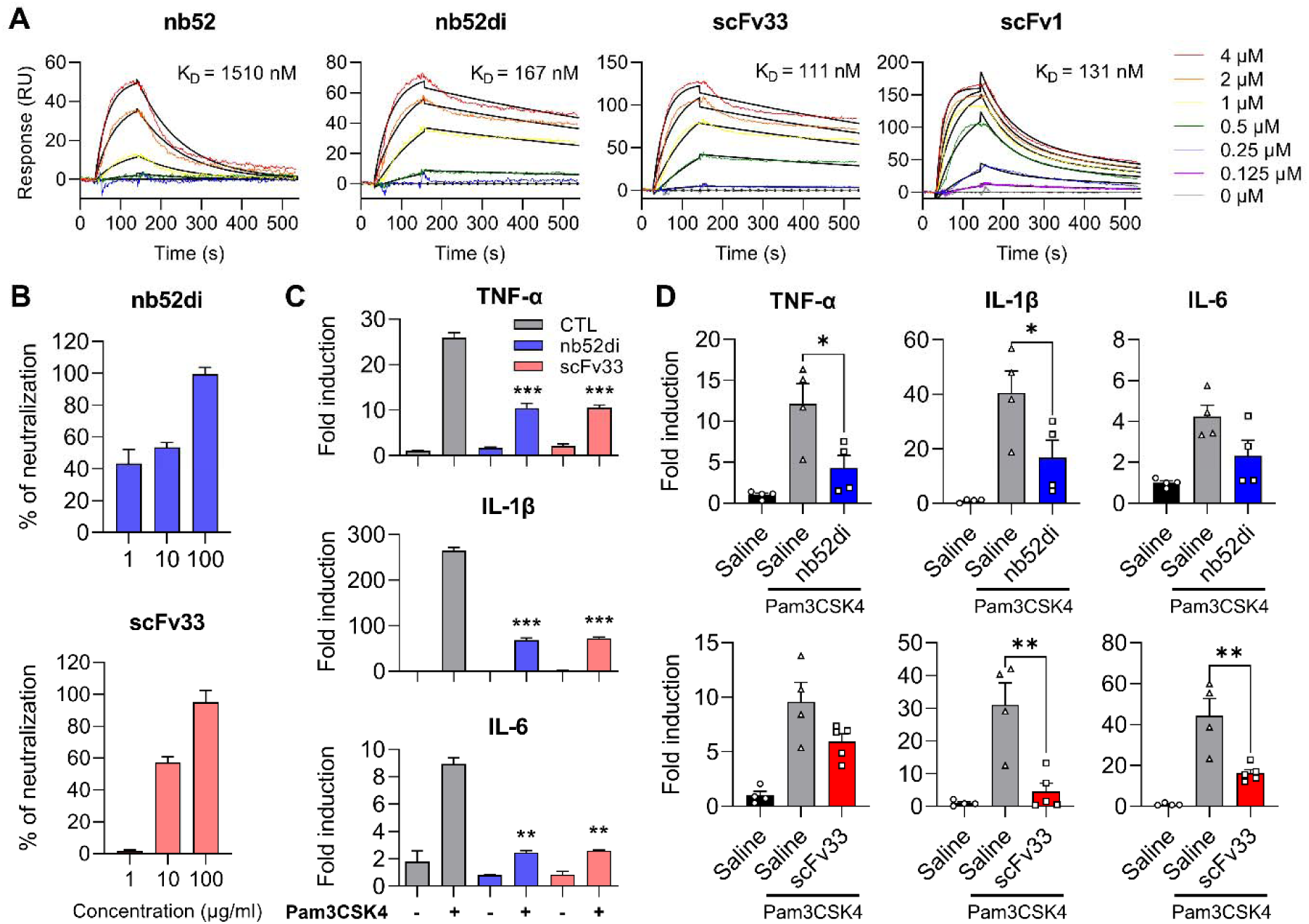
High-affinity anti-TLR2 nanobodies and scFvs inhibit NF-κB activation and downstream inflammation. (**A**) SPR analysis showing dose-dependent binding of nb52, nb52 dimer (nb52di), scFv33, and scFv1 to immobilized recombinant TLR2. Equilibrium dissociation constants (KD) are indicated. **(B)** TLR2/NF-κB signaling inhibition assessed by a SEAP reporter assay. HEK-Blue™ hTLR2 cells were pretreated with nb52di or scFv33 for 1 hour and stimulated with Pam3CSK4 (0.5 ng/mL) for 48 hours. SEAP activity was quantified as OD655, and neutralization was expressed as a percentage relative to the Pam3CSK4-only controls. Data represent the mean ± SEM of 2–3 technical replicates. **(C)** qRT-PCR analysis of proinflammatory cytokine expression in BV2 cells. Cells were pretreated with nb52di (50 μg/mL) or scFv33 (30 μg/mL) for 1 hour and stimulated with Pam3CSK4 (50 ng/mL) for 3 hours. Expression levels of TNF-α, IL-1β, and IL-6 are shown relative to unstimulated controls. Data represent the mean ± SEM of 2 technical replicates. Statistical analysis was performed using two-way ANOVA followed by Tukey’s post hoc test (**p < 0.01, ***p < 0.001, ****p < 0.0001 vs. Pam3CSK4 group). **(D)** qRT-PCR analysis of proinflammatory cytokine expression in mouse spinal cords (L4–L6). Mice received a single intrathecal injection of Pam3CSK4 (10 ng) with and without the nb52 dimer or scFv33 (2 μg) in 10 μL of saline. Each dot represents an individual mouse. Data represent the mean ± SEM. Statistical analysis was performed using one-way ANOVA with Tukey’s post hoc test (*p < 0.05, **p < 0.01 vs. Pam3CSK4+saline group).

To assess functional blocking activity for TLR2, we performed a secreted embryonic alkaline phosphatase (SEAP) reporter assay using HEK-Blue™ human TLR2 cells, which secrete SEAPs upon TLR2-mediated NF-κB activation. SEAP activity in the culture supernatant was measured as OD_655_. Pre-treatment with the nb52 dimer or scFv33 suppressed Pam3CSK4-induced SEAP activity in a dose-dependent manner, with 100 μg achieving near-complete (∼100%) inhibition (Figure 2B).

Given their ability to inhibit TLR2 signaling, an important question was whether these antibodies could suppress downstream inflammatory responses. To answer that question, we pretreated BV2 and THP-1 cells with the antibodies and then stimulated them with Pam3CSK4. The expression levels of proinflammatory cytokines were quantified by qRT-PCR. In both cell lines, the nanobodies and scFvs significantly reduced TNF-α, IL-1β, and IL-6 expression, compared with the saline-treated control (Figures 2C and S4). Notably, in BV2 cells, the nb52 dimer and scFv33 significantly suppressed TNF-α, IL-1β, and IL-6 expression by approximately 60%, 70%, and 75%, respectively (Figure 2C). To extend those findings *in vivo*, we treated mice with intrathecal Pam3CSK4 with and without the nb52 dimer or scFv33. After 6 hours, the lumbar spinal cord (L4–L6) was harvested for qRT-PCR analysis. The nb52 dimer significantly reduced TNF-α and IL-1β by 60–70%; scFv33 suppressed IL-1β and IL-6 by ∼86% and 63%, respectively (Figure 2D).

Taken together, these results demonstrate that the selected anti-TLR2 antibodies—especially the nb52 dimer and scFv33—bind TLR2 with high affinity and specificity and effectively inhibit TLR2-mediated inflammatory responses both *in vitro* and *in vivo*, supporting their therapeutic potential for neuroinflammatory diseases.

### In vivo analgesic efficacy of anti-TLR2 nanobodies and scFvs in neuropathic pain models

We next evaluated the *in vivo* analgesic efficacy of the selected anti-TLR2 nanobody and scFv using the spinal nerve transection (SNT) model of neuropathic pain. Mechanical allodynia was assessed using electronic von Frey testing, and the mice received a single intrathecal injection of one antibody (2 μg) on day 3 post-surgery. All antibody-treated groups exhibited increased withdrawal thresholds, compared with the saline group, starting from day 1 post-injection (Figure 3B). Both the nb52 monomer and dimer elevated withdrawal thresholds to ∼3.5 g, markedly higher than the saline group (2.4–2.8 g), during the first two weeks, with the effect waning after day 14–18 post-injection. scFv33 induced a more moderate increase (3.0–3.5 g) but maintained that level through day 21, with statistical significance still observed on day 18.

**Figure 3.**
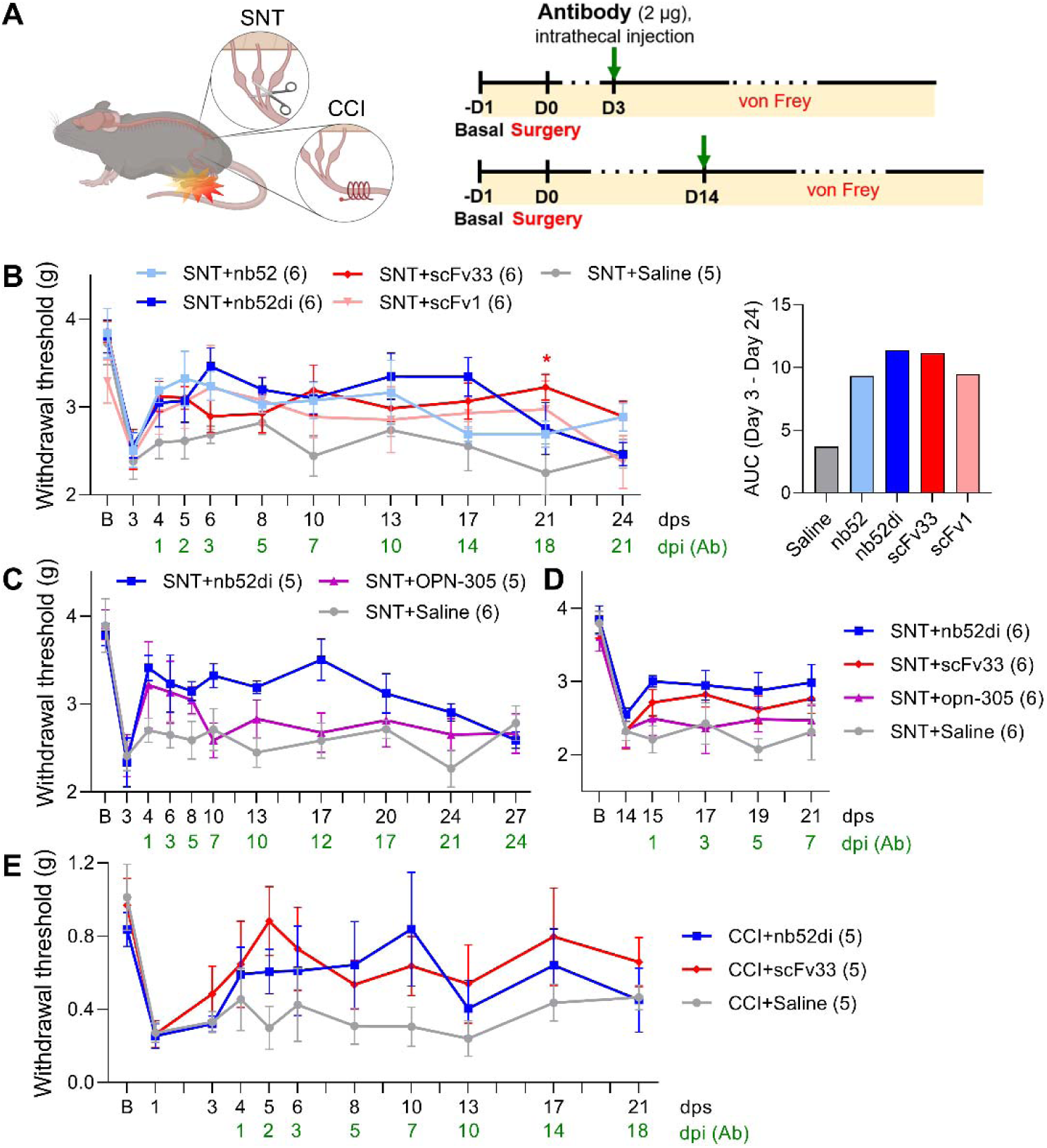
Analgesic efficacy of the anti-TLR2 nanobodies and scFvs in mouse neuropathic pain models. **(A)** Schematic illustration of the SNT and CCI models of nerve injury–induced neuropathic pain and the experimental timeline for antibody treatment and behavioral testing. **(B–D)** Mechanical withdrawal thresholds measured by electronic von Frey in mice subjected to SNT, following a single administration of the antibody (Ab, 2 μg) on day 3 **(B,C)** or day 14 **(D)** post-surgery. **(C)** includes comparison with OPN-305 as a conventional anti-TLR2 antibody. **(E)** Mechanical withdrawal thresholds assessed using von Frey filaments in mice subjected to CCI, following a single administration of the Ab on day 3 post-surgery. Data represent the mean ± SEM. Numbers in parentheses indicate the number of animals per group. Statistical significance was determined by two-way ANOVA with Tukey’s post hoc test (*p < 0.05 vs. SNT + saline). dps, days post-surgery; dpi, days post-injection.

Based on their superior binding affinity to TLR2 (Figure 2A) and *in vivo* analgesic effects in the SNT model (Figure 3B), the nb52 dimer and scFv33 were selected for further evaluation. To compare them with a clinically tested anti-TLR2 antibody (OPN-305),^23^ mice received the same amount of nb52 dimer or OPN-305 on day 3 post-surgery. Both treatments increased the withdrawal thresholds to ∼3.5 g, but the nb52 dimer sustained analgesia for nearly three weeks, whereas OPN-305’s effect declined sharply after day 7 (Figure 3C).

Given that treatment often begins in the chronic pain stage, we evaluated delayed therapeutic efficacy. Those mice received a single intrathecal injection of the nb52 dimer, scFv33, or OPN-305 on day 14 post-surgery. Both the nb52 dimer and scFv33 increased the withdrawal thresholds (∼3.0 g) from day 1 post-injection and maintained their effects through day 7, whereas OPN-305 showed no noticeable analgesia (Figure 3D).

To further evaluate their efficacy, we tested the same antibodies in the chronic constriction injury (CCI) model, another widely used model of peripheral neuropathic pain. Following CCI surgery, mice were injected on day 3 and assessed using von Frey filaments. Both the nb52 dimer and scFv33 modestly increased withdrawal thresholds (0.7–0.9 g) compared with saline, and that trend persisted for up to two weeks, although without statistical significance (Figure 3E).

The *in vivo* safety profile of the selected antibodies was evaluated by repeated intrathecal administration, once per week for 3 weeks. No significant changes were observed in body weight or open field locomotor activity, compared with control animals (Figure S5), indicating no apparent toxicity or adverse effects.

Taken together, these results demonstrate that the selected anti-TLR2 antibodies—particularly the nb52 dimer and scFv33—exert robust and sustained analgesic effects in mouse models of neuropathic pain, including when they are administered at a delayed therapeutic time point, and they outperformed the clinically tested antibody OPN-305.

### In vitro and in vivo evaluation of PLGA nanoparticles to deliver anti-TLR2 antibody genes

Although the recombinant anti-TLR2 antibodies alone exhibited significant analgesic effects, the chronicity and persistence of neuropathic pain highlights the need for therapeutic strategies with prolonged efficacy. Therefore, we sought to incorporate a gene therapy approach using PLGA NPs to achieve sustained *in vivo* expression of the antibodies. We first constructed mammalian expression vectors encoding the nb52 dimer or scFv33; the vectors incorporated a PelBκ signal peptide for secretion,^24,25^ a C-terminal His-tag, and an IRES-EGFP cassette for transfection monitoring (Figure 4A and S6A). Transient transfection of the vectors into HEK293T cells confirmed robust EGFP fluorescence and intracellular His-tag immunostaining, indicating successful expression (Figure S6B). Western blot analyses further demonstrated that both antibodies were secreted into the culture medium (Figure S6C).

**Figure 4.**
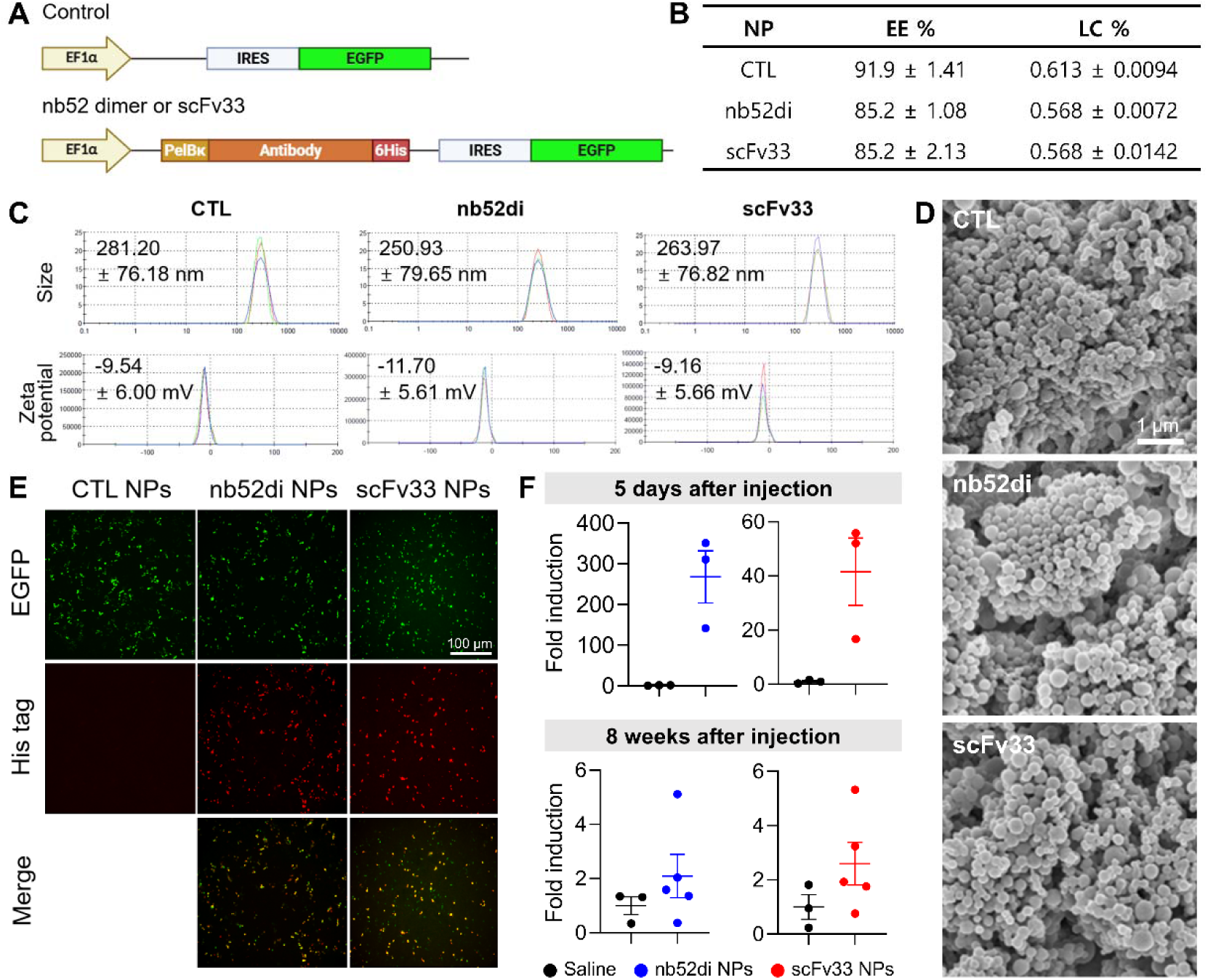
Characterization and *in vivo* expression of PLGA nanoparticles encapsulating anti-TLR2 antibody genes. **(A)** Schematic of the mammalian expression vector encoding the nb52 dimer or scFv33, under the control of the EF1α promoter. **(B)** Encapsulation efficiency (EE) and loading capacity (LC) of PLGA NPs encapsulating control (CTL), nb52 dimer (nb52di), and scFv33 plasmids. **(C)** Particle size and zeta potential of the respective nanoparticles, measured using a Zetasizer. **(D)** Scanning electron microscopy images of PLGA NPs loaded with control (CTL), nb52 dimer (nb52di), or scFv33 antibody genes. **(E)** Immunofluorescence analysis of HEK293T cells transfected with CTL, nb52di, or scFv33 NPs, 48 h post-transfection. **(F)** mRNA expression of the nb52 dimer and scFv33, as analyzed by qRT-PCR, following an intrathecal injection of nb52di or scFv33 PLGA NPs (200 μg). Fold induction was calculated relative to the saline control. Each dot represents an individual mouse. Data represent the mean ± SEM.

To deliver the antibody genes *in vivo*, we used a double emulsion (water-in-oil-in-water, W_1_/O/W_2_) method to formulate PLGA NPs that encapsulated each plasmid.^26,27^ The resulting NPs exhibited high encapsulation efficiency (≥ 85%) and loading capacity of 0.57–0.61% (Figure 4B; see Equation 1 and 2 in Method details). All NPs showed particle sizes between 250 and 280 nm and moderately negative zeta potentials (–9 to –12 mV) (Figure 4C), which are favorable for uptake by phagocytic cells such as microglia.^24,28^ We confirmed the uniform spherical morphology of all PLGA NPs by scanning electron microscopy (Figure 4D). The biocompatibility of PLGA NPs was confirmed by MTS assays in primary glial cultures that showed no cytotoxicity, i.e., viability maintained above 95%, across a range of concentrations (0–400 μg/mL) (Figure S7).

To determine whether the PLGA NPs could successfully mediate anti-TLR2 antibody gene delivery, initial validation was performed *in vitro* using HEK293T cells. Cells were transfected with PLGA NPs encapsulating nb52 dimer or scFv33 pDNA. After 48 hours, strong EGFP expression and co-localized His-tag signals were observed, confirming successful NP-mediated gene transfer and expression of the antibodies (Figure 4E). For *in vivo* validation, mice received an intrathecal injection of 200 μg of NPs, and the spinal lumbar segment (L4–L6) was collected at 5 days and 8 weeks post-injection. qRT-PCR showed strong induction of nb52 dimer and scFv33 mRNA at 5 days and detectable expression persisting up to 8 weeks, confirming long-term gene expression from a single NP administration (Figure 4F).

Collectively, these results demonstrate that PLGA NPs were robustly synthesized with consistent physicochemical properties, that they effectively enabled gene delivery of anti-TLR2 nanobody and scFv constructs both *in vitro* and *in vivo*, and that they notably provided long-term production of functional therapeutic proteins within the target tissues.

### Enhanced and long-lasting analgesic effect of PLGA nanoparticle–mediated anti-TLR2 antibody gene delivery in pain models

After confirming the successful delivery and expression of the anti-TLR2 nanobody and scFv via PLGA NPs, we next assessed whether that approach could provide therapeutic efficacy superior to the recombinant antibody injection. In the SNT model, a single intrathecal injection of nb52 dimer or scFv33 NPs significantly alleviated mechanical allodynia, compared with saline-treated controls, and the effects persisted for more than 8 weeks (Figure 5B).

**Figure 5.**
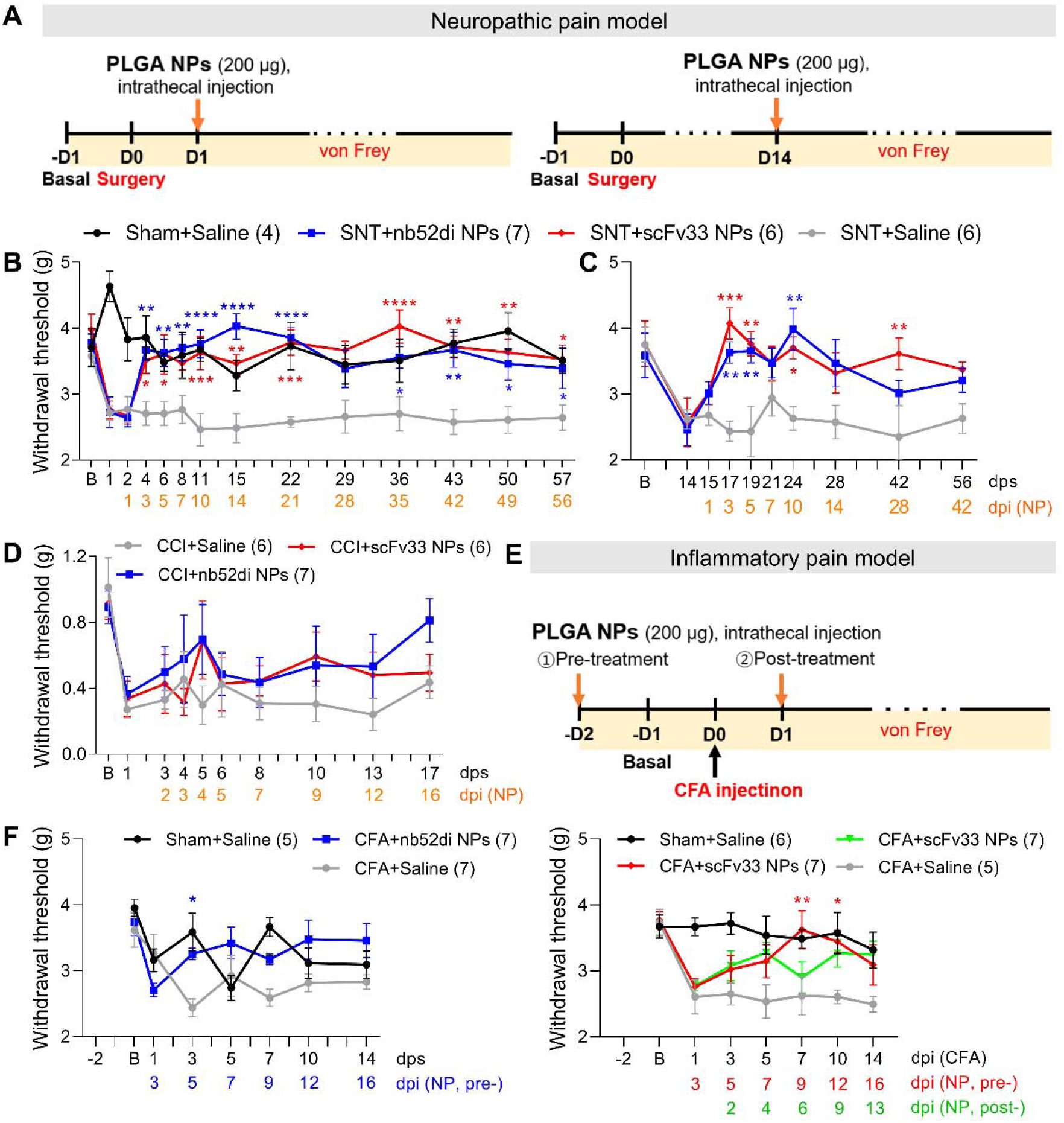
Enhanced and sustained analgesia by PLGA nanoparticle–mediated delivery of anti-TLR2 antibody genes. (**A,E**) Schematic illustration of the experimental timelines for the neuropathic **(A)** and inflammatory **(E)** pain models. **(B,C)** Mechanical withdrawal thresholds in the SNT model were measured using the electronic von Frey test following an intrathecal injection of PLGA NPs. **(D)** Mechanical allodynia thresholds in the CCI model were assessed using the von Frey filament test. **(F)** Mechanical withdrawal thresholds in the CFA-induced inflammatory pain model (using pre– and/or post-treatment paradigms) were measured with the electronic von Frey test. All data represent the mean ± SEM. Numbers in parentheses indicate the number of animals per group. Statistical significance was determined by two-way ANOVA with Tukey’s post hoc test (*p < 0.05, **p < 0.01, ***p < 0.001, ****p < 0.0001 vs. saline group subjected to surgery or CFA injection). dps, days post-surgery; dpi, days post-injection.

Furthermore, we evaluated their analgesic efficacy at delayed time points following nerve injury to increase their clinical relevance. An administration of nb52 dimer or scFv33 NPs on day 7 or 14 post-surgery significantly reversed hypersensitivity in SNT mice, compared with saline-treated controls, and the effects lasted for 6–8 weeks (Figures S8A and 5C). Similarly, scFv33 NPs administered on day 21 post-surgery reversed mechanical hypersensitivity significantly on day 28 post-injection, although the effect was slower and somewhat reduced compared with the day 7 and 14 injections (Figure S8B). Nonetheless, the analgesic effect lasted for up to four weeks after injection. Similar effects were observed in the CCI model, though the magnitude of analgesia was less pronounced (Figure 5D), likely reflecting model-specific immune responses; CCI involves milder and more diffuse inflammatory signaling than SNT.

Following the demonstration of robust analgesic effects with a 200 μg dose of nb52 dimer or scFv33 NPs in an SNT mouse model, we investigated the analgesic efficacy of lower doses. Although onset appeared slightly delayed, even 50 μg and 20 μg of nb52 dimer NPs produced significant analgesic effects (Figure S8C).

To broaden the therapeutic relevance, we next assessed their efficacy in the CFA-induced inflammatory pain model. Pre-treatment with nb52di or scFv33 NPs (two days before CFA injection) significantly prevented the development of mechanical hypersensitivity, with elevated withdrawal thresholds maintained for at least 14 days (Figure 5F). ScFv33 NPs administered two days after CFA injection also alleviated hypersensitivity, although the therapeutic effect was more modest than pre-treatment (Figure 5F, right).

To exclude the possibility that the NPs had non-specific effects, control PLGA NPs encapsulating non-coding plasmids were intrathecally administered to mice subjected to SNT or CFA (Figure S8D,E). The control NP-treated mice showed no significant difference in withdrawal thresholds, compared with the saline controls, confirming that the analgesic effects observed with the nb52 dimer and scFv33 NPs were specifically mediated by anti-TLR2 antibody gene expression.

Together, these findings demonstrate that PLGA NP–mediated delivery of anti-TLR2 antibody genes provides sustained analgesia in both neuropathic and inflammatory pain, with flexible efficacy across doses and time points and durability superior to the recombinant antibody protein injection.

### Suppression of spinal neuroinflammation and glial activation by PLGA NP–mediated anti-TLR2 antibody gene delivery

To investigate the mechanism underlying the analgesic effects of anti-TLR2 antibody gene delivery, we analyzed inflammatory responses in the spinal cords of SNT mice following an intrathecal administration of nb52 dimer or scFv33 NPs on day 1 post-surgery. Lumbar spinal cord segments (L4–L6) were collected on day 4 and day 7 post-surgery for qRT-PCR analysis. Significant downregulation of proinflammatory cytokines (TNF-α, IL-1β, IL-6) was observed on day 7 in both treatment groups (Figure 6A,B). A supplementary analysis further showed that the nb52di NPs significantly reduced the expression of the oxidative stress mediators Nox2 and iNOS on day 4, suggesting attenuation of early upstream inflammatory signaling. TLR2 expression exhibited a decreasing trend on day 7 (Figure S9).

**Figure 6.**
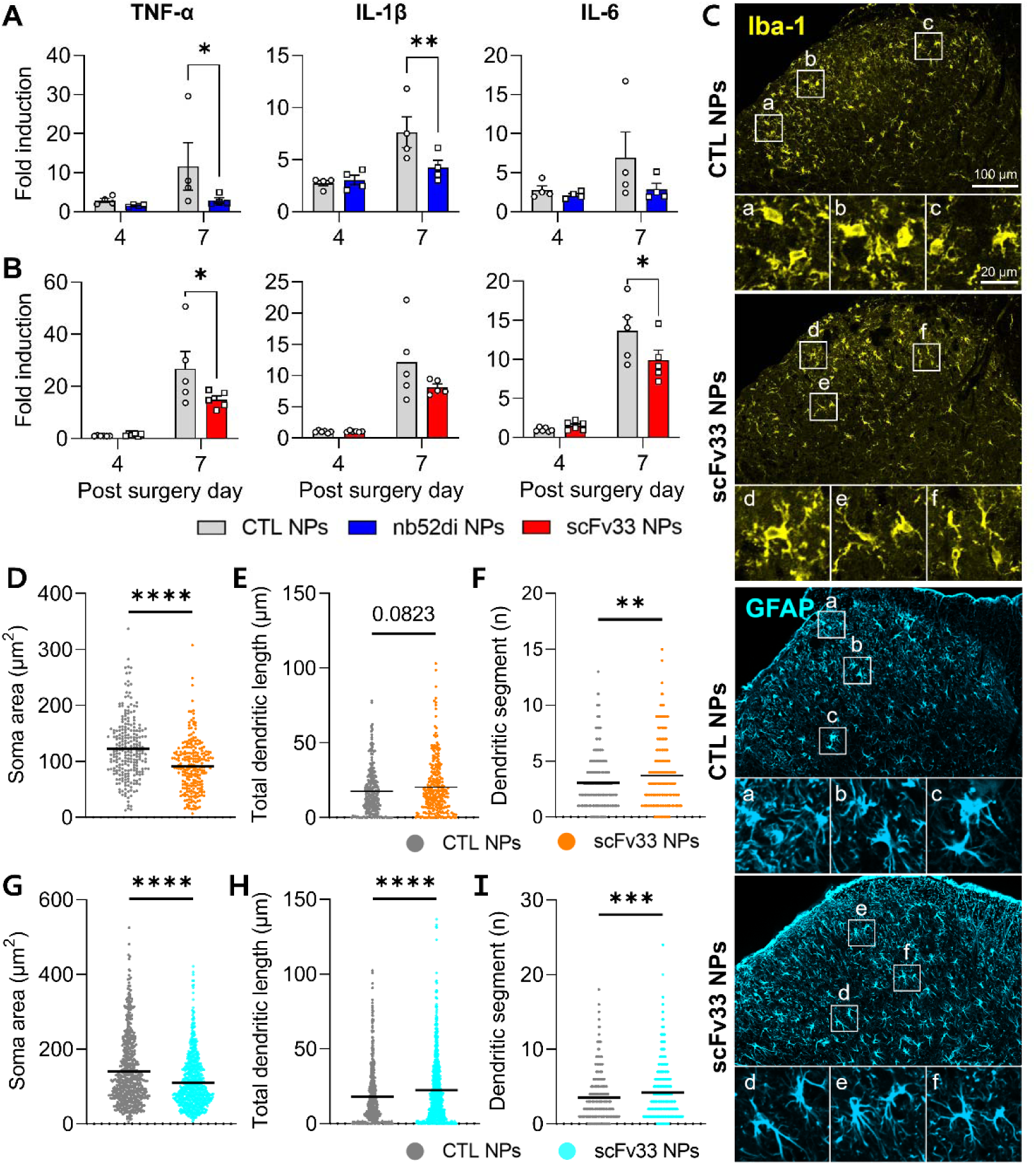
Inhibition of neuroinflammatory signaling and glial activation by PLGA nanoparticle–mediated anti-TLR2 antibody gene delivery. SNT-subjected mice were intrathecally injected with nb52 dimer (nb52di), scFv33, or control (CTL) NPs (200 μg) on day 1 post-surgery. Spinal cord segments (L4–L6) were harvested for qRT-PCR and immunohistochemistry analyses. **(A,B)** mRNA expression of proinflammatory cytokines was analyzed by qRT-PCR on days 4 and 7 post-surgery. The expression levels of TNF-α, IL-1β, and IL-6 are shown relative to the saline-injected sham group. Each dot represents an individual mouse. Data represent the mean ± SEM. Statistical analysis was performed using two-way ANOVA followed by Tukey’s post hoc test (*p < 0.05, **p < 0.01 vs. CTL NP group). **(C)** Representative immunohistochemistry images collected on day 7 post-surgery. **(D–F)** Morphological analyses of individual Iba-1^+^ microglia. **(G–I)** Morphological analyses of individual GFAP^+^ astrocytes. Each dot represents an individual cell. Data represent the mean ± SEM. Statistical significance was determined using unpaired t-tests (*p < 0.05, **p < 0.01, ***p < 0.001, ****p < 0.0001 vs. control NPs).

We next examined glial cell activation by an immunohistochemical analysis of Iba-1 and GFAP in spinal cord sections harvested on day 7. Although no significant differences were observed in Iba-1 or GFAP fluorescence intensity, coverage, or cell density (Figure S10), a morphological analysis of individual glial cells revealed significant changes following scFv33 NP treatment. Microglia from treated mice exhibited reduced soma area and increased dendritic segment number, indicative of a less activated phenotype than the controls (Figure 6D–F). Astrocytes also showed reduced soma size and increased dendritic length and complexity (Figure 6G–I).

Together, these findings indicate that PLGA NP–mediated delivery of the anti-TLR2 nanobody and scFv genes effectively suppresses SNT-induced neuroinflammation and induces a morphological shift in glial cells toward a less reactive state that contributes to the observed analgesic effects.

## DISCUSSION

TLR2 is a key pattern recognition receptor in the innate immune system that recognizes a wide array of PAMPs and DAMPs.^29,30^ Upon activation, TLR2 triggers downstream signaling pathways that culminate in NF-κB activation and the subsequent transcription of proinflammatory cytokines, chemokines, and immune mediators. In the CNS, microglia express high levels of TLR2 and respond to both endogenous and exogenous stimuli by initiating neuroinflammatory responses.^31^

In neuropathic pain models, peripheral nerve injury induces sustained microglial activation in the spinal cord, accompanied by upregulation of TLR2 expression and increased production of proinflammatory cytokines, including TNF-α, IL-1β, and IL-6. Those cytokines contribute to central sensitization and the maintenance of chronic pain states.^32^

In this study, we developed recombinant anti-TLR2 antibodies, including the nb52 dimer and scFv33, to specifically inhibit TLR2 signaling pathways and modulate the microglial activation implicated in the pathogenesis of neuropathic pain. Using synthetic nanobody and human-derived scFv libraries, high-affinity candidates were identified through phage display and ELISA-based screening methods, and the selected nb52 dimer and scFv33 were the lead molecules (Figures 1 and 2).

Functional assays demonstrated that both the nb52 dimer and scFv33 effectively suppressed Pam3CSK4-induced NF-κB activation in TLR2-overexpressing HEK293T cells, as well as the mRNA expression of proinflammatory cytokines both *in vitro* and *in vivo* (Figure 2). Furthermore, in the SNT model of neuropathic pain, an administration of recombinant nb52 dimer or scFv33 proteins significantly alleviated mechanical allodynia, compared with saline-treated controls, and showed greater efficacy than OPN-305, a clinically evaluated anti-TLR2 antibody (Figure 3). These results indicate that the recombinant anti-TLR2 nanobody and scFv not only inhibit TLR2-mediated inflammatory signaling, but also provide analgesic effects superior to existing IgG antibody therapeutics.

Although nanobodies and scFvs offer advantages over conventional full-length antibodies, antibody treatments remain limited in chronic disease contexts that require sustained molecular intervention. Therefore, we used PLGA NPs for *in vivo* gene delivery of the anti-TLR2 antibody constructs. *In vivo* studies in the mouse SNT model of neuropathic pain demonstrated that a single intrathecal administration of nb52 dimer or scFv33 NPs significantly alleviated mechanical allodynia (Figures 5 and S8). Notably, the analgesic effects persisted more than 2.5 times longer than those observed following direct antibody injection. This prolonged effect was accompanied by reduced activation of microglia and astrocytes, as well as the downregulation of proinflammatory cytokine expression in the spinal cord (Figure 6), suggesting the suppression of TLR2-mediated neuroinflammation, one of the key mechanisms underlying central sensitization. These findings highlight the therapeutic potential of PLGA NP–mediated gene delivery for the long-term management of chronic pain.

Compared with the SNT model, anti-TLR2 antibody treatment yielded relatively modest analgesic effects in the CCI model (Figures 3E and 5D). This difference likely stems from the distinct temporal and molecular characteristics of the two nerve injury paradigms. SNT induces an immediate and localized axonal injury near the dorsal root ganglia that leads to rapid TLR2-dependent microglial activation and transient upregulation of proinflammatory mediators such as NOX2, TNF-α, and IL-1β. In this context, TLR2 acts as a key upstream regulator of early neuroinflammation, and its inhibition effectively reduces mechanical allodynia.^11,12,33^ In contrast, CCI elicits a slower and more complex inflammatory response. Spinal *Tlr2* expression begins to rise only after day 2 post-surgery and continues to increase through at least day 14, in parallel with *Tlr4*.^34^ Furthermore, single-cell RNA sequencing has shown that by post-surgery day 7, microglia upregulate a large complement/chemokine module (*C4b*, *Cfb*, *Ccl5*, *Cxcl10*) that is largely absent in SNT. This indicates a broader and more redundant innate immune network in CCI.^35^ As a result, selective TLR2 blockade might be insufficient to fully suppress neuroinflammation in this model due to the compensatory activation of parallel pathways. This underscores the importance of identifying disease-specific mechanisms, optimizing therapeutic timing, and ensuring precise molecular targeting.

PLGA NP–mediated gene delivery of anti-TLR2 antibodies produced robust analgesic effects in the SNT model, not only during the acute phase but also when administered at delayed time points—7, 14, and 21 days post-injury (Figures 5C and S8A,B). However, the therapeutic onset was progressively delayed and the magnitude of behavioral recovery diminished slightly as the injection was postponed. These findings suggest that TLR2-mediated inflammation is most active during the early phase of nerve injury but continues to contribute to chronic pain maintenance. This is supported by previous studies showing sustained microglial activation beyond the acute phase of neuropathic injury. In SNT models, spinal Iba-1 levels remain elevated at 7–14 days post-injury.^36,37^ Similar trends have been observed in other models, such as spinal nerve ligation injury, in which microglial markers remain elevated at 21 days post-injury.^38^ Together, these findings indicate that although microglial activation peaks during the early phase of nerve injury, it persists at subacute levels for extended periods. This chronic, low-grade neuroinflammation could contribute to sustained nociceptive sensitization and pain recurrence. Therefore, therapeutic strategies targeting microglial signaling—such as TLR2 blockade—remain mechanistically relevant even during the chronic phase and might offer therapeutic benefit beyond the initial injury window.

Several challenges remain. First, mechanistic profiling was confined to NF-κB and glial cell activation; comprehensive phospho-proteomics or bulk RNA sequencing are needed to determine whether a TLR2 blockade also suppresses other TLR2-related signaling pathways, such as MAPK (p38/JNK) and IRAK4 cascades,^39,40^ and define the contributions of astrocytes or neurons that also express TLR2.^41,42^ Second, although no overt toxicity or motor deficits were observed, long-term safety—especially infection risk under sustained TLR2 inhibition and an accumulation of PLGA-derived metabolites—must be tested in aged or immunocompromised hosts. This highlights the need for reversible control systems of gene therapy (e.g., chemical-inducible promoters or CRISPR-off circuits) that enable the regulation of therapeutic antibody expression in a time– and dose-controlled manner *in vivo*.^43–45^

In summary, we developed recombinant nanobodies and scFvs targeting microglial TLR2, a key driver of inflammatory pathology in injury-driven pain hypersensitivity. Furthermore, we introduced PLGA NP–based gene therapy as a delivery platform for anti-TLR2 antibody genes to obtain durable efficacy. Both the nb52 dimer and scFv33 were successfully expressed in mouse spinal cords via PLGA NP-mediated gene delivery, sustaining the expression for months, tempering glial reactivity, and rescuing behavior in various pain models. These results establish a preclinical proof-of-concept that PLGA NP-mediated receptor-specific gene immunotherapy is a feasible strategy for long-term treatment of chronic pain.

## MATERIALS AND METHODS

### Library construction

A synthetic nanobody library was constructed based on a humanized single-domain antibody (hs2dAb) scaffold derived from the sdAbD10 framework, which was previously optimized for intracellular stability and solubility^22^. Randomized CDRs were designed based on natural VHH diversity and introduced using trinucleotide-directed synthesis, with fixed lengths of 7 amino acids for CDR1 and CDR2, and variable lengths of 8 to 18 amino acids for CDR3. The resulting library fragments were PCR-amplified. A human scFv phage-display library (HuScL-7; Creative Biolabs, Shirley, NY, USA) was purchased.

Synthesized oligomers containing PelBκ signal peptide sequence were inserted into the pADL23c phagemid vector (Antibody Design Labs, San Diego, CA, USA) between EcoRI and NcoI (Enzynomics, Daejeon, Korea). The antibody library was digested with HindIII (Enzynomics, Daejeon, Korea), and NotI (New England Biolabs, Ipswich, MA, USA), and then ligated into the modified pADL23c phagemid vector containing a His-tag, a myc-tag, and an ampicillin-resistance gene for positive clone selection using T4 DNA ligase (Thermo Fisher Scientific, Waltham, MA, USA).

For phage display library preparation, the ligation products were electroporated into *E. coli* TG1 cells (Antibody Design Labs, San Diego, CA, USA) using the Gene Pulser Xcell™ Electroporation System (Bio-Rad Laboratories, Hercules, CA, USA) at 2500 V, 25 μF, and 200 Ω.

Following electroporation, the transformed *E. coli* TG1 cells were cultured in 250 mL baffled flasks containing 20 mL of LB medium (2×YT, 2% glucose, 100 μg/mL ampicillin) at 37 °C with shaking (180 rpm) for 2–3 h until the optical density reached OD_600_ = 0.5. The cultures were then infected with CM13 helper phage (Antibody Design Labs, San Diego, CA, USA) at a 10:1 multiplicity of infection (2 × 10 pfu/mL) and incubated for an additional 1 h under the same conditions. After infection, the cultures were centrifuged at 6,000 rpm for 5 min at 4 °C, the supernatant was discarded, and the cell pellet was resuspended in 20 mL of LB medium (2×YT, 100 μg/mL ampicillin, 50 μg/mL kanamycin). The cultures were incubated overnight at 30 °C with shaking (180 rpm) to allow phage particle production. The next day, the cultures were centrifuged at 10,000 rpm for 10 min at 4 °C, and the supernatant was collected and filtered through a 0.45 μm syringe filter.

To precipitate the phage particles, 4 mL of PEG/NaCl precipitation buffer (PPB; 20% PEG 8000, 2.5 M NaCl) was added, and the mixture was incubated for at least 2 h on a shaker at 4 °C. The phage particles were pelleted by centrifugation at 12,000 rpm for 30 min at 4 °C, resuspended in 800 μL of PBS by gentle pipetting, and further incubated with an additional 200 μL of PPB for 15 min at 4 °C on a rotator. A second centrifugation was then performed at 12,000 rpm for 15–30 min at 4 °C, after which the supernatant was discarded and the final phage pellet was gently resuspended in 200 μL of PBS. The phage titer was determined by infection of E. coli TG1 cells with serial dilutions of the preparation.

### Phage display biopanning

Biopanning of the synthetic nanobody library was conducted using recombinant protein immobilized on 96-well ELISA plates (Nunc MaxiSorp, Thermo Fisher Scientific, Roskilde, Denmark). Each well was coated with 50 μL of recombinant human TLR2 protein (40 μg/mL; MyBioSource, San Diego, CA, USA) in coating buffer (0.1 M NaHCO_3_, pH 8.3) and incubated overnight at 4 °C with gentle shaking. The next day, the wells were blocked with 3% skim milk in PBS for 1 h at room temperature with agitation at 110 rpm. After one wash with PBST (PBS containing 0.1% Tween-20), 1 × 10¹¹ phage particles in 100 μL of PBS containing 0.1% (v/v) Tween-20 (PBST) were added per well and incubated for 1 h at room temperature with gentle shaking.

Following incubation, the wells were washed five times (first round) or ten times (second to fourth rounds) with 0.1% PBST. Each wash step consisted of a 2-minute incubation followed by inversion and blotting on paper towels to remove unbound phages. Bound phages were eluted by adding 100 μL per well of elution buffer (0.2 M Glycine-HCl, pH 2.2, containing 0.1% BSA) and incubating for 15 min at room temperature with shaking. Eluted phages were vigorously pipetted 10 times, transferred to 1.5 mL tubes, and neutralized with 15 μL of Tris-HCl buffer (1 M, pH 9.1) per well. The total neutralized eluate from two wells was pooled and used for subsequent phage titration and amplification.

### ELISA selection with phages

To obtain crude protein, E. coli SS320 (Antibody Design Labs, San Diego, CA, USA) was prepared in 10 mL of LB/tetracycline (25 μg/mL) at 37 °C until OD_600_ reached 0.5. A 500 μL of the E. coli SS320 suspension was infected with 10 μL of phage supernatant and incubated for 10 min at room temperature. The infected cells were plated onto LB agar containing 100 μg/mL ampicillin and incubated overnight at 37 °C.

The next day, a single colony was inoculated into 10 mL of LB medium (2×YT, 100 μg/mL ampicillin) and cultured at 37 °C until OD_600_ reached 0.5. Protein expression was induced with 0.2 mM IPTG, followed by overnight incubation at 30 °C with shaking. For ELISA coating, 50 μL of induced bacterial culture (diluted to a final antigen concentration of 2–4 μg/mL) was added to each well of a 96-well plate (Corning, Cat. No. 9017) and incubated overnight at 4 °C with gentle shaking.

After removing the coating solution, the wells were blocked with 3% (w/v) skim milk in 0.1% PBST at room temperature for 1 h. To prepare crude protein extracts, induced cultures were centrifuged at 6,000 rpm for 5 min at 4 °C, and the cell pellets were resuspended in 500 μL of TES buffer (200 mM Tris-HCl, 1 mM EDTA, pH 8.0, 20% sucrose). After incubation on ice for 15 min, the cells were centrifuged at 13,000 rpm for 2 min, resuspended in 500 μL of ice-cold distilled water, and incubated again on ice for 15 min. The crude protein-containing supernatant was collected after a final centrifugation at 13,000 rpm for 2 min at 4 °C.

Blocking buffer was removed from the ELISA plates, and each well was incubated with 100 μL of a 3:1 mixture of blocking buffer and crude protein solution for 1 h at room temperature with gentle agitation. Plates were washed five times with 0.1% PBST, followed by incubation with 50 μL of either anti-Myc-HRP antibody (for nanobody detection; Santa Cruz Biotechnology, 1:500) or anti-E tag-HRP antibody (for scFv detection; Abcam, 1:5,000) for 1 h at room temperature. After washing, 50 μL of TMB substrate solution (Thermo Fisher Scientific, Waltham, MA, USA) was added to each well and incubated for approximately 15 min at room temperature. The reaction was stopped by adding 50 μL of stop solution, and absorbance was measured at 450 nm using an EMax® Plus Microplate Reader (Molecular Devices, San Jose, CA, USA).

### ELISA selection with crude protein

To prepare crude protein, E. coli SS320 was prepared in 10 mL of LB/tetracycline (25 μg/mL) at 37 °C until OD_600_ reached 0.5. A 500 μL of the E. coli SS320 suspension was infected with 10 μL of phage supernatant and incubated for 10 min at room temperature. The infected cells were plated onto LB agar containing 100 μg/mL ampicillin and incubated overnight at 37 °C. The next day, a single colony was inoculated into 10 mL of LB medium (2×YT, 100 μg/mL ampicillin) and cultured at 37 °C until OD_600_ reached 0.5. Protein expression was induced with 0.2 mM IPTG, followed by overnight incubation at 30 °C with shaking. The induced cultures were centrifuged at 6,000 rpm for 5 min at 4 °C, and the cell pellets were resuspended in 500 μL of TES_2_ buffer (200 mM Tris-HCl, 1 mM EDTA, pH 8.0, 20% sucrose). After incubation on ice for 15 min, cells were centrifuged at 13,000 rpm for 2 min, resuspended in 500 μL of ice-cold distilled water, and incubated again on ice for 15 min. The crude protein-containing supernatant was collected after a final centrifugation at 13,000 rpm for 2 min at 4 °C.

For ELISA coating, 50 μL of recombinant human TLR2 protein (2 μg/mL) was added to each well of a 96-well plate (Corning Costar, Corning Inc., Corning, NY, USA) and incubated overnight at 4 °C with gentle shaking. After removing the coating solution, the wells were blocked with 3% skim milk in 0.1% PBST at room temperature for 1 h. Blocking buffer was removed from the ELISA plates, and each well was incubated with 100 μL of a 3:1 mixture of blocking buffer and crude protein solution for 1 h at room temperature with gentle agitation. Plates were washed five times with 0.1% PBST, followed by incubation with 50 μL of either anti-Myc-HRP antibody for nanobody detection (Santa Cruz Biotechnology, Dallas, TX, USA; dilution 1:500) or anti-E tag-HRP antibody (Abcam, Cambridge, UK; dilution 1:5,000) for 1 h at RT. After washing, 50 μL of TMB solution was added to each well and incubated for approximately 15 min at room temperature. The reaction was stopped by adding 50 μL of stop solution, and absorbance was measured at 450 nm using a microplate reader.

Following ELISA screening using crude protein, the corresponding phage clones were selected for PCR amplification and DNA sequencing. A single colony from each positive clone was used as a template for colony PCR using a nanobody/scFv-specific primer set. Amplified DNA fragments were purified and subjected to Sanger sequencing. Based on DNA sequence analysis, primary candidate clones were selected for further validation.

### Antibody purification

For protein expression, a single colony of E. coli harboring the nanobody or scFv construct was inoculated into 5 mL of LB medium (100 μg/mL ampicillin) and cultured overnight at 37 °C. The following day, a 1:100 diluted pre-culture was transferred into 200 mL of fresh LB/ampicillin medium in a 500 mL baffled flask and grown at 37 °C until the OD_600_ reached approximately 0.5. Protein expression was induced by adding IPTG to a final concentration of 0.1 mM, followed by incubation with shaking at 37 °C for 5 h.

The induced culture was harvested by centrifugation and subjected to osmotic lysis using TES buffer (200 mM Tris-HCl, 1 mM EDTA, pH 8.0, 20% sucrose) and ice-cold distilled water supplemented with protease and phosphatase inhibitor cocktail (Thermo Fisher Scientific, Waltham, MA, USA). Cell lysates were clarified by centrifugation and filtered for affinity purification.

Purification of His-tagged recombinant proteins was carried out using Ni-NTA His•Bind® Resin (Merck, Darmstadt, Germany) packed into gravity-flow columns. After column equilibration with 1× binding buffer (300 mM NaCl, 50 mM sodium phosphate, 10 mM imidazole, pH 7.4), clarified lysates were loaded. The column was then washed with wash buffer (300 mM NaCl, 50 mM sodium phosphate, 20 mM imidazole, pH 7.4), and proteins were eluted using elution buffer (300 mM NaCl, 50 mM sodium phosphate, and 250 mM imidazole, pH 7.4). Eluates were pooled and either stored at –20 °C or used immediately.

To reduce endotoxin contamination, purified proteins were treated with 1% Triton X-114 and subjected to multiple cycles of phase separation. The endotoxin-depleted supernatant was concentrated and buffer-exchanged into PBS using centrifugal filter units (10 kDa MWCO; MilliporeSigma, Burlington, MA, USA).

Protein concentration was determined using a BCA assay (Thermo Fisher Scientific, Waltham, MA, USA), and purity was assessed by SDS-PAGE and western blot using anti-His antibodies (Santa Cruz Biotechnology, Dallas, TX, USA).

### Western Blot

To evaluate the expression and secretion of the recombinant anti-TLR2 antibody following pDNA transfection, both cell lysates and culture supernatants were collected for immunoblotting.

For cell lysate preparation, transfected cells were lysed with RIPA buffer containing protease inhibitors (1 mM), sodium orthovanadate (1 mM), and sodium fluoride (1 mM). After incubation on ice for 10 min, the cell suspension was collected, centrifuged at 12,000 rpm for 10 min at 4 °C, and the supernatant was transferred to a fresh tube.

For analysis of secreted protein, 400 μL of culture medium was mixed with 1.6 mL of ice-cold acetone (final 4:1 ratio), vortexed briefly, and incubated at –20 °C for 1 h. Samples were then centrifuged at 12,000 rpm for 15 min at 4 °C. The supernatant was discarded, and the protein pellet was air-dried for 10 min. The pellet was resuspended in 200 μL of lysis buffer, vortexed, and briefly sonicated. After an additional centrifugation step (12,000 rpm, 15 min, 4 °C), the final supernatant was collected while avoiding any insoluble debris.

Equal amounts of protein samples were mixed with 5× SDS sample buffer in lysis buffer and boiled. Proteins were separated on 12% SDS-PAGE gels and transferred onto 0.22 μm nitrocellulose membranes (GE Healthcare, Chicago, IL, USA) at 300 mA for 1 h. Membranes were blocked in 5% skim milk in TBS with 0.1% Tween-20 for 1 h at room temperature, followed by overnight incubation at 4 °C with anti-His antibody (1:1,000; Santa Cruz Biotechnology, Dallas, TX, USA). The next day, membranes were washed, incubated with anti-mouse IgG-HRP (Abfrontier, Seoul, Republic of Korea), and developed using WESTSAVE™ Gold ECL Solution (Abfrontier, Seoul, Republic of Korea).

### Surface Plasmon Resonance (SPR) analysis

Surface plasmon resonance (SPR) analysis was performed using a gold sensor chip-based SPR system (IMSPR-ProX, ICluebio, Seoul, Korea) to evaluate the binding kinetics between recombinant TLR2 and purified antibody candidates.

For TLR2 protein coating on a sensor chip, the surface was first stabilized and a baseline was established by flowing the running buffer (PBST, 0.05% Tween-20) for 2 min. The COOH-Au chip (Icluebio, Ansan, Republic of Korea) was then activated using the Amine Coupling Kit (Icluebio, Ansan, Republic of Korea), which contains NHS solution, EDC, and ethanolamine-HCl quenching buffer. After pre-injection of borate buffer (30 s at 50 μL/min; Polysciences, Warrington, PA, USA), a freshly prepared 1:1 mixture of EDC (0.4 M, prepared in acetate buffer provided in the kit) and NHS solution (0.1 M, supplied in the kit) was injected (30 s at 50 μL/min) to activate the carboxyl groups on the chip surface. Recombinant human TLR2 protein was diluted in acetate buffer (pH 4.0; 15 μg in 300 μL) and injected to covalently immobilize the ligand onto the sensor surface (20 min at 10 μL/min). The surface was subsequently blocked with 100 μg/mL BSA in acetate buffer (pH 4.0) and quenched with ethanolamine-HCl quenching buffer (5 min at 50 μL/min). Unbound materials between each step were removed by running buffer (30 s at 50 μL/min). Chip regeneration was performed using glycine-HCl buffer (pH 2.5) between analyte injections.

Antibody samples were prepared in PBST (0.05% Tween-20) and injected at concentrations ranging from 0 to 500 nM in two-fold serial dilutions (60 s at 50 μL/min), followed by a dissociation phase for 5 min and chip regeneration for 3 min. All samples were run in duplicate. Sensorgrams (CH1–CH2) were collected in real-time and analyzed using integrated kinetic analysis software. Kinetic parameters, including the association rate constant (k_a_), dissociation rate constant (k_d_), and equilibrium dissociation constant (K_D_), were calculated using a 1:1 Langmuir binding model.

### Secreted embryonic alkaline phosphatase (SEAP) reporter assay

To assess the inhibitory effects of antibodies on TLR2 signaling, SEAP reporter assays were performed using HEK-Blue™ hTLR2 cells (InvivoGen, San Diego, CA, USA), which stably express human TLR2 and a SEAP reporter gene under the control of an NF-κB-inducible promoter.

HEK-Blue™ hTLR2 cells (InvivoGen, San Diego, CA, USA) were seeded into 96-well plates at a density of 1 × 10 cells per well and incubated overnight at 37 °C in a humidified incubator with 5% CO_2_. The following day, the culture medium was replaced with 160 μL of HEK-Blue™ Detection medium (InvivoGen), and cells were treated with 20 μL of recombinant antibody solutions in PBS. After 1 h of antibody pretreatment, cells were stimulated with 0.5 ng/mL Pam3CSK4 (Tocris Bioscience, Bristol, UK) and incubated for 48 h. SEAP activity in the supernatant was assessed by measuring OD at 655 nm using a microplate reader (Molecular Devices, San Jose, CA, USA).

### Animals

All animal procedures were conducted with approval from the Institutional Animal Care and Use Committee (IACUC) of Seoul National University. Male C57BL/6J mice (8–10 weeks old) were purchased from Daehan Biolink (DBL, Eumsung, Korea). 5xFAD transgenic mice on a C57BL/6 background [B6SJL-Tg(APPSwFlLon,PSEN1M146LL286V)6799Vas/Mmjax; RRID:MMRRC_034840-JAX] were provided by the Korea Research Institute of Bioscience and Biotechnology (KRIBB, Daejeon, Republic of Korea). They were housed under specific pathogen-free (SPF) conditions at 22–24 °C and 55% relative humidity, with a 12-h light/dark cycle. Food and water were provided ad libitum. All experiments were carried out in accordance with the guidelines of the International Association for the Study of Pain (IASP).

### Quantitative real-time PCR (qRT-PCR)

Total RNA was extracted from BV2 microglial cells, THP-1 monocytes, or mouse lumbar spinal cord segments (L4–L6) using TRIzol™ Reagent (Invitrogen, Thermo Fisher Scientific, Waltham, MA, USA), and cDNA was synthesized according to the manufacturer’s protocol. Quantitative real-time PCR was performed using 40 ng of cDNA per reaction with SYBR™ Green PCR Master Mix (Thermo Fisher Scientific, Waltham, MA, USA) on a StepOnePlus™ Real-Time PCR System (Applied Biosystems, Foster City, CA, USA).

The amplification program was as follows: initial denaturation at 95 °C for 15 min; two cycles of 94 °C for 15 s and 49 °C for 15 s; followed by 32 cycles of 94 °C for 15 s, 62 °C for 15 s, and 74 °C for 15 s with signal acquisition during the extension step; and a final step at 84 °C for 10 s and 88 °C for 15 s.

The following PCR primer sequences were used: GAPDH forward, 5′-AGT ATG ACT CCA CTC ACG GCA A-3′; GAPDH reverse, 5′-TCT CGC TCC TGG AAG ATG GT-3′; TNF-α forward, 5′-GGC TCT TCT GGA TCT TGG TG-3′; TNF-α reverse, 5′-TTT CAT GGC TGC TGT GAG TC-3′; IL-1β forward, 5′-TTG TGG CTG TGG AGA AGC TGT-3′; IL-1β reverse, 5′-AAC GTC ACA CAC CAG CAG GTT-3′; IL-6 forward, 5′-TCC ATC CAG TTG CCT TCT TGG-3′; IL-6 reverse, 5′-CCA CGA TTT CCC AGA GAA CAT G-3′. Gene expression levels were normalized to mouse GAPDH and expressed as fold changes relative to the control group, calculated using the 2-ΔΔCt method, as previously described.^46^

### Intrathecal injection

For delivery of the recombinant antibodies or PLGA NPs to the mouse spinal cord or brain, mice were anesthetized with isoflurane in an O_2_ carrier (induction 2% and maintenance 1.5%). A total volume of 10 μL was administered using a 10 μL or 50 μL Hamilton syringe (Hamilton Company, Reno, NV, USA) with a 30-gauge needle.

For spinal delivery, the solution was injected intrathecally into the L5–L6 intervertebral space while the lower back was gently flexed by hand. Successful administration was confirmed by a brief tail flick response. For delivery to the brain, the needle was inserted percutaneously into the cisterna magna with the head fixed in a flexed position by hand. A total volume of 10 μL was slowly injected over 30 s. The needle was left in place for additional 10 s before removal, and then direct pressure was applied to the puncture site.

### Pain models

#### Spinal Nerve Transection (SNT) Model

The SNT model was established as previously described.^12^ Mice were anesthetized with isoflurane delivered in oxygen (induction 2% and maintenance 1.5%), and placed in the prone position. A small skin incision was made to expose the lumbar vertebrae, and blunt dissection was performed to visualize the L4 and L5 spinal nerves. The L6 transverse process was carefully removed to expose the L5 spinal nerve, which was then completely transected using fine surgical scissors. After surgery, the incision was closed with surgical staples, and mice were allowed to recover in their home cages. Sham-operated mice underwent the same procedure except that the L5 spinal nerve was left intact. All surgical steps were performed under aseptic conditions to prevent infection and minimize inflammatory responses.

#### Chronic Constriction Injury (CCI) Model

The CCI model was established as previously described.^47^ Under anesthesia with isoflurane in oxygen (induction 2% and maintenance 1.5%), the right sciatic nerve was exposed at the mid-thigh level through a small incision. The nerve was loosely tied at four sites, approximately 1 mm apart, using 6-0 chromic gut sutures (Ailee Co., Busan, Korea). After ligation, the overlying muscles and skin were closed with the same suture material. Sham surgeries followed the same procedure except that the sciatic nerve was not ligated. Mice were returned to their home cages for recovery. All surgical steps were performed under aseptic conditions.

#### Complete Freund’s Adjuvant (CFA) Model

The CFA model was established as previously described.^48^ Mice received a subcutaneous injection of 10 μL CFA emulsion (mixed 1:1 with sterile saline; Sigma-Aldrich, St. Louis, MO, USA) into the plantar surface of the left hind paw. Control mice received sterile saline only. After injection, all animals were returned to their home cages for recovery. All surgical steps were performed under aseptic conditions.

### Immunocytochemistry (ICC)

Cells cultured on plates were washed with sterile 0.1 M PBS and fixed with 2% paraformaldehyde. Fixed cells were permeabilized with 0.1% Triton X-100 in 0.1 M PBS, then incubated with 0.1 M PBS containing 5% normal goat serum (Jackson ImmunoResearch, West Grove, PA, USA), 2% BSA, and 0.1% Triton X-100 for 1 h. Samples were then incubated overnight at 4 °C with anti-His-Tag antibody (Santa Cruz Biotechnology, Dallas, TX, USA). Following three washes in 0.1 M PBS, samples were incubated for 2 h at room temperature with secondary antibodies conjugated to Cy3 (1:200; Jackson ImmunoResearch, West Grove, PA, USA), and washed five times in 0.1 M PBS. Fluorescent images were obtained using a fluorescence microscope (DP72 / Olympus BX51; Olympus, Tokyo, Japan) or a confocal microscope (LSM800; Carl Zeiss).

### PLGA nanoparticle synthesis

PLGA NPs encapsulating pDNA were synthesized based on a previously described double emulsion solvent evaporation method.^26^ Briefly, 100 μg of pDNA dissolved in 200 μL of TE buffer (pH 8.0) was added dropwise into 1 mL of dichloromethane (DCM; Samchun Chemicals, Seoul, Republic of Korea) containing 20 mg of PLGA (Resomer® RG 502H; Sigma-Aldrich, St. Louis, MO, USA). The mixture was emulsified by probe sonication (Vibra-Cell™ VCX 130; Sonics, Newtown, CT, USA) at 40 W for 1 min to generate a primary water-in-oil (W1/O) emulsion. This primary emulsion was then transferred into 2 mL of 1% polyvinyl alcohol (Alfa Aesar, Ward Hill, MA, USA) and sonicated for 2 min to form a water-in-oil-in-water (W1/O/W2) double emulsion. The resulting mixture was diluted with 6 mL of 1% polyvinyl alcohol solution containing 12.5 mM CaCl_2_ and stirred at room temperature for 3 h in a fume hood to allow DCM evaporation. Synthesized NPs were collected by centrifugation at 15,000 × g for 15 min at 4 °C (Hanil Scientific Inc., Daejeon, Korea), washed twice with deionized water, and lyophilized for storage.

### Nanoparticle characterization

#### Measurement of Particle Size, Surface Charge, and Morphology

The hydrodynamic diameter and zeta potential of the NPs were measured by dynamic light scattering using a Zetasizer Nano-ZS instrument (Malvern Instruments, Malvern, UK). Scanning electron micrographs of the NPs were obtained using a field-emission scanning electron microscope (Carl Zeiss Microscopy GmbH, Jena, Germany).

#### Encapsulation Efficiency (EE) and Loading Capacity (LC)

Following the sonication and solvent evaporation steps during nanoparticle preparation, PLGA NPs were washed to remove unencapsulated pDNA and excess polymer. The nanoparticles were pelleted by centrifugation, and the unencapsulated DNA remaining in the supernatant was quantified. EE and LC were calculated according to the following equations^49,50^:

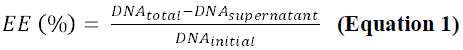

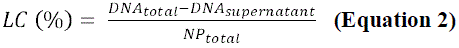

### Primary mixed glia culture

Primary mixed glial cultures were prepared from the whole brains of postnatal day 1–2 neonatal mice. Brain tissues were dissociated and cultured in high-glucose DMEM (4.5 g/L) supplemented with 10% fetal bovine serum (FBS), 10 mM HEPES, 2 mM L-glutamine, 1× Anti-Anti, and 1× non-essential amino acids (all from Gibco, Thermo Fisher Scientific, Waltham, MA, USA). Glial cultures were incubated at 37 °C with 5% CO_2_, and the culture medium was refreshed every 5 days. After 15 days in vitro, microglial cells were detached using 0.05% Trypsin–EDTA (Gibco, Thermo Fisher Scientific, Waltham, MA, USA) and seeded on 96-well plates for the MTS assay.

### MTS assay

Cell viability was assessed using an MTS-based colorimetric assay. Mixed glial cells were seeded into 96-well plates at a density of 0.5 × 10^4^ cells per well and incubated for 24 h. Cells were then treated with PLGA NPs for 24 h. After treatment, 20 μL of MTS reagent (Promega, Madison, WI, USA) was added directly to each well, and the plate was incubated at 37 °C for 1 h. The MTS tetrazolium compound was reduced by metabolically active cells to generate a water-soluble formazan product. Absorbance was measured at 490 nm using a microplate reader (Molecular Devices, San Jose, CA, USA). Cell viability was calculated by comparing absorbance values of treated wells to those of untreated control wells.

### Immunohistochemistry (IHC)

Mice were transcardially perfused with 0.1 M phosphate buffer (pH 7.4), followed by 4% paraformaldehyde. The L4–L6 spinal cord was dissected and post-fixed overnight in the same fixative at 4 °C. Tissues were then cryoprotected in 30% sucrose for at least 48 h, embedded and sectioned coronally at 16 μm thickness using a cryostat (CM1860; Leica, Wetzlar, Germany).

Tissue sections were blocked for 1 h at room temperature in 0.1 M PBS containing 5% normal goat serum (Jackson ImmunoResearch, West Grove, PA, USA), 2% BSA, and 0.2% Triton X-100. Samples were then incubated overnight at 4 °C with anti-Iba1 (1:1,000; FUJIFILM Wako, Osaka, Japan) and anti-GFAP (1:1,000; Abcam, Cambridge, UK). Following three washes in 0.1 M PBS, sections were incubated for 2 h at room temperature with secondary antibodies conjugated to FITC and Cy3 (1:200; Jackson ImmunoResearch, USA). Following five washes in 0.1 M PBS, stained sections were mounted on glass slides using Vectashield mounting medium (Vector Laboratories, Burlingame, CA, USA). Fluorescent images were obtained using a confocal microscope (LSM800; Carl Zeiss). Glial cell activation was analyzed using IMARIS (version 9.8.0, Oxford Instruments, Abingdon, UK).

### Behavioral test

#### Pain assessment

Mechanical allodynia was assessed using an electronic von Frey system (dynamic plantar aesthesiometer; Ugo Basile, Milan, Italy), as previously described.^51^ Mice were individually placed in transparent chambers on an elevated wire mesh platform, and after acclimatization, a progressively increasing force was applied to the plantar surface of the ipsilateral hind paw. The force was ramped linearly up to 5 g over 10 s and was then held constant at 5 g for an additional 30 s. The paw withdrawal threshold was defined as the force at which the animal withdrew its paw. Baseline measurements were obtained prior to surgery, and all behavioral testing was performed under blinded conditions.

In the CCI model, mechanical sensitivity was assessed using a set of manual von Frey filaments (0.02–4 g; Stoelting, Wood Dale, IL, USA) and the up–down method94. Positive responses were defined as sharp paw withdrawal, flinching, or licking following filament application. The 50% withdrawal threshold was calculated using the Dixon up–down algorithm based on the animal’s response pattern.

## QUANTIFICATION AND STATISTICAL ANALYSIS

All statistical analyses were performed using GraphPad Prism version 7.0 for Windows (GraphPad Software Inc., La Jolla, CA, USA). Data are presented as the mean ± standard error of the mean (SEM). Comparisons among multiple groups were assessed using one-way or two-way analysis of variance (ANOVA) followed by Tukey’s post hoc test, unless otherwise specified in the figure legends. Differences were considered statistically significant when the p-value was less than 0.05.

Some of the figures in this study were created using BioRender.com, a web-based science illustration tool. Plasmid maps were generated using SnapGene (from Insightful Science; available at snapgene.com).

## ACKNOWLEDGEMENTS

This research was supported by GliaCellTech Inc. (Project No. 860-20230106).

## AUTHOR CONTRIBUTIONS

Subeen Lee conducted the investigation, performed formal analysis and validation, prepared visualizations, and wrote the original draft of the manuscript. Jaekyung Jeon and Hyunji Lee contributed to the investigation, formal analysis, validation, and visualization. Ellane Eda Barcelona supported the validation experiments. Jinpyo Hong contributed to the conceptualization and methodology of the study and provided supervision. Sung Joong Lee conceived and supervised the project, acquired funding and resources, managed the project administration, and revised the manuscript.

