## Supplementary figures for "Development of Recombinant Anti-TLR2 Antibodies and PLGA Nanoparticle-based Gene Therapy for the Treatment of Neuropathic Pain"

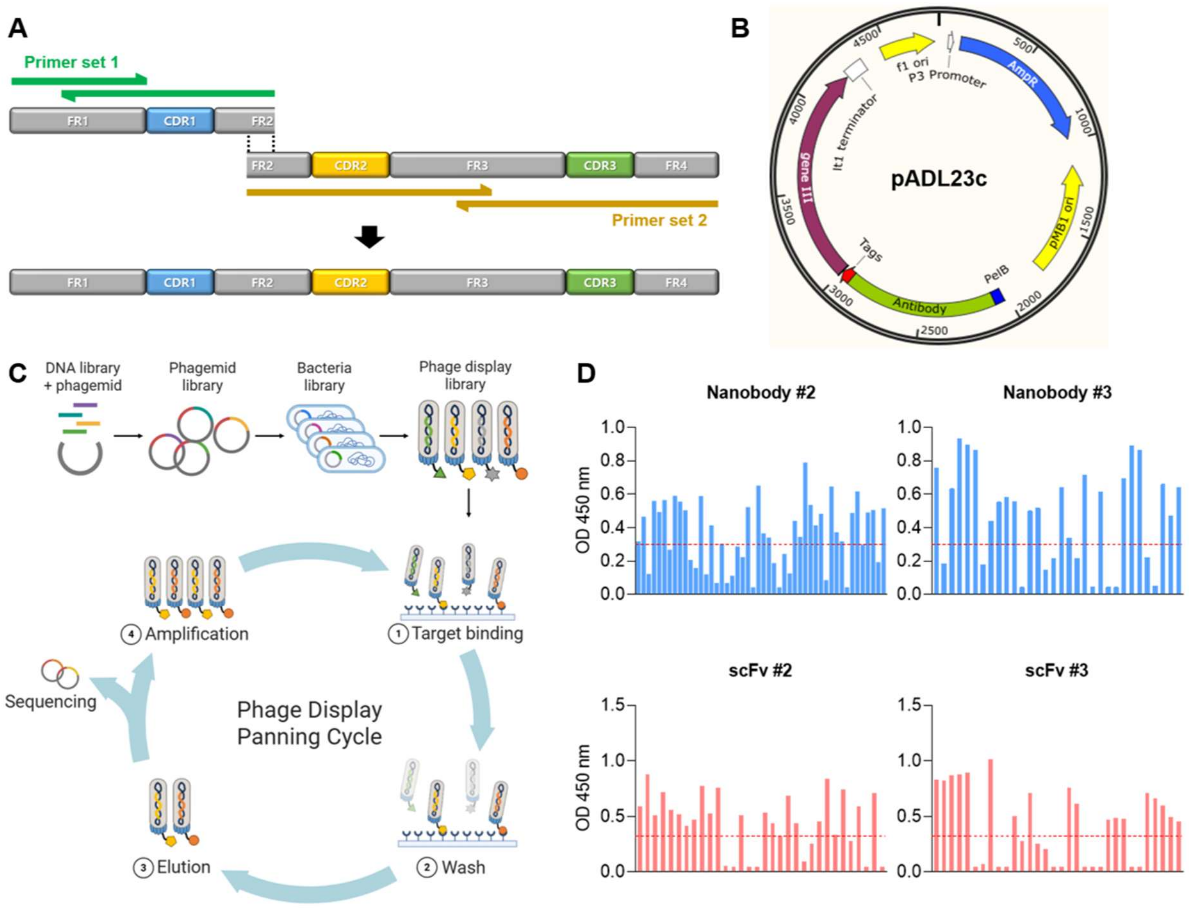


**Supplementary Figure 1. Construction and screening of phage-displayed nanobody and scFv libraries targeting TLR2. (A)** Schematic representation of the two-step PCR strategy used to assemble full-length nanobody genes with randomized CDRs. **(B)** Map of the pADL23c phagemid vector used to clone the nanobody constructs. The vector includes an amber stop codon between the antibody and p**III** sequences. **(C)** Workflow of the phage display biopanning used to enrich anti-TLR2 phage clones through repeated rounds of selection. **(D)** Additional ELISA screening data for individual phage-displayed clones obtained and amplified after the final round of biopanning.

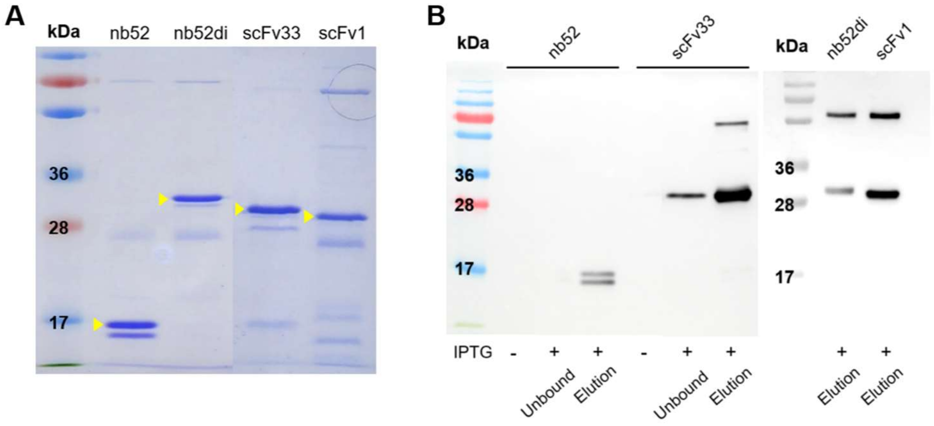


**Supplementary Figure 2. Production and purification of the selected recombinant anti-TLR2 antibodies in *E. coli.* (A)** SDS-PAGE analysis of the anti-TLR2 nanobody and scFv candidates (nb52 monomer and dimer, scFv33, and scFv1) expressed in SS320 *E. coli* upon Isopropyl β-D-1-thiogalactopyranoside (IPTG) induction and purified using Ni-NTA affinity chromatography. Distinct bands were observed at the expected molecular weights (indicated by yellow arrowheads). **(B)** Immunoblot analysis using an anti-His antibody confirmed the expression and purification of the recombinant anti-TLR2 antibody clones. His-tagged antibody bands were detected in IPTG-induced (+) lysates and Ni-NTA eluates. No bands were observed in uninduced (–) samples. Unbound lanes represent wash flow-through fractions before elution.

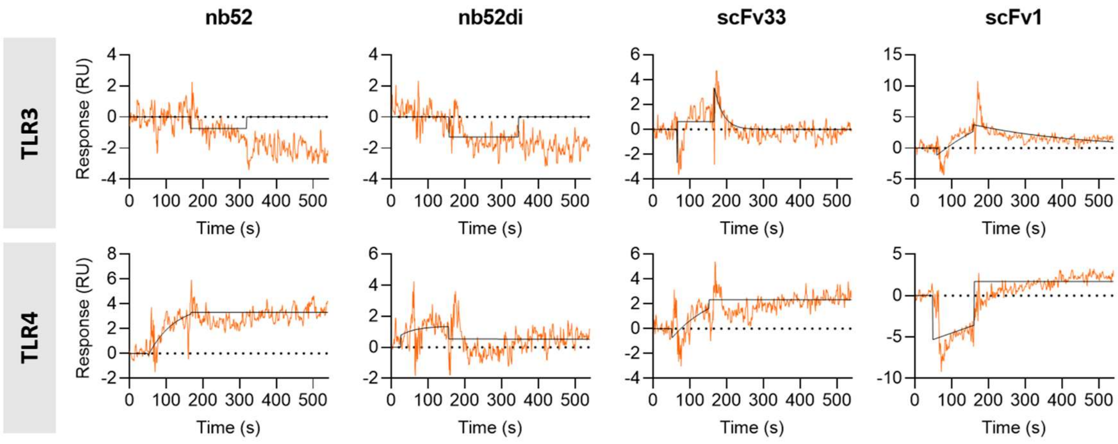


**Supplementary Figure 3. Binding activity of the selected anti-TLR2 recombinant antibodies measured by SPR analysis against TLR3 and TLR4.** Surface plasmon resonance (SPR) analysis of nb52, nb52 dimer (nb52di), scFv33, and scFv1 was performed at 250 nM against immobilized recombinant human TLR3 and TLR4 proteins. No measurable binding was observed for any clone, indicating high specificity of the selected antibodies for TLR2.

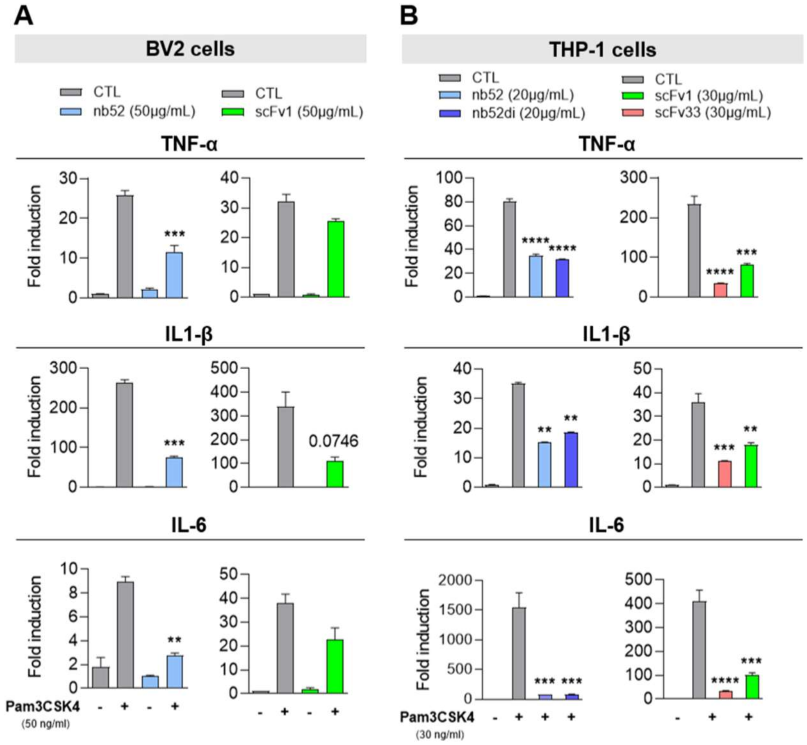


**Supplementary Figure 4. The selected anti-TLR2 antibodies suppress Pam3CSK4-induced inflammatory responses in BV2 and THP-1 cells. (A,B)** qRT-PCR analysis of proinflammatory cytokine expression in BV2 cells **(A)** and THP-1 cells **(B)**. Cells were pretreated with nb52, nb52di, scFv33, or scFv1 for 1 hour and stimulated with Pam3CSK4 (50 ng/mL) for 3 hours. Expression levels of TNF-α, IL-1β, and IL-6 are shown relative to unstimulated controls. Data represent the mean ± SEM of 2 technical replicates. Statistical analysis was performed using two-way ANOVA followed by Tukey's post hoc test (**$p < 0.01$, ***$p < 0.001$, ****$p < 0.0001$ vs. Pam3CSK4 group).

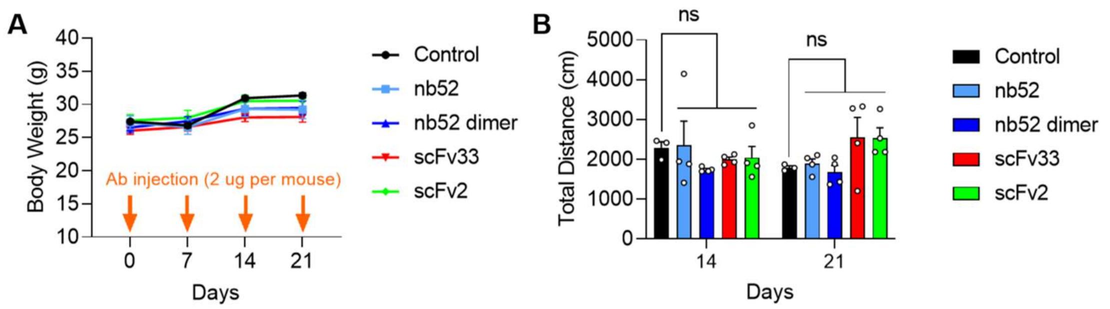


**Supplementary Figure 5. Safety assessment of anti-TLR2 nanobodies and scFvs following repeated intrathecal administration. (A,B)** Mice received repeated weekly intrathecal injections of the indicated nanobodies and scFvs (2 µg) to assess potential toxicity. **(A)** Body weight was monitored for 21 days. **(B)** Locomotor activity was evaluated in the open field test by measuring the total distance traveled on days 14 and 21. Each dot represents an individual mouse. Data represent the mean ± SEM. Statistical analysis was performed using one-way ANOVA with Tukey’s post hoc test.

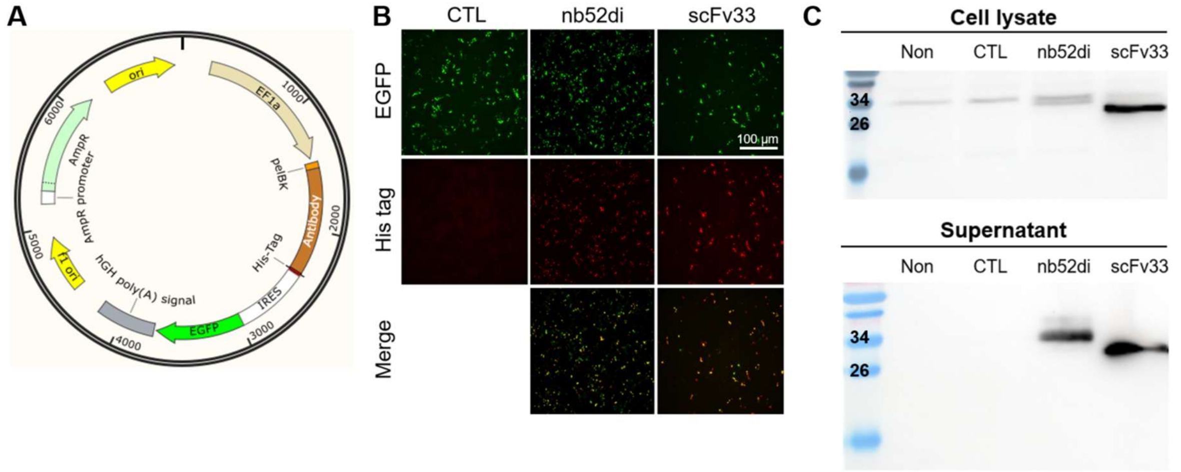


**Supplementary Figure 6. Expression and secretion of an anti-TLR2 nanobody and scFv in HEK293T cells following vector transfection. (A)** Schematic map of the mammalian expression vector encoding the nb52 dimer or scFv33 antibody under the control of the EF1α promoter. Constructs contain an N-terminal PelBκ signal peptide for secretion and a C-terminal 6×His tag for detection. An internal ribosome entry site followed by EGFP was included as a fluorescent reporter. **(B)** 48 h after transfection with control (CTL), nb52 dimer (nb52di), or scFv33 expression vectors. Merged images show co-localization of EGFP and His-tag signals. **(C)** Western blot analysis of cell lysates and culture supernatants from transfected HEK293T cells, probed with an anti-His antibody.

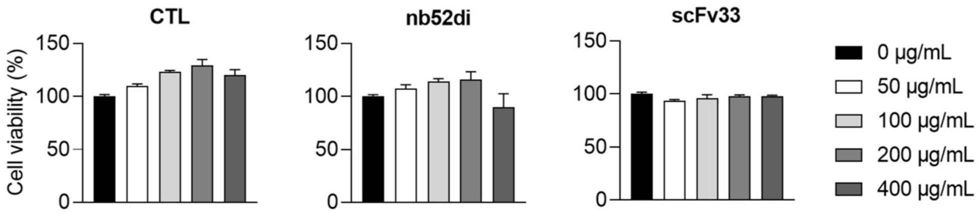


**Supplementary Figure 7. Cytotoxicity assessment of PLGA nanoparticles in primary mixed glial cultures.** Cell viability of primary mixed glial cultures treated with increasing concentrations (0–400 μg/mL) of PLGA NPs for 24 hours, measured by the MTS assay. Data are expressed as the percent viability relative to untreated controls (0 μg/mL) and are shown as the mean ± SEM of 3 technical replicates.

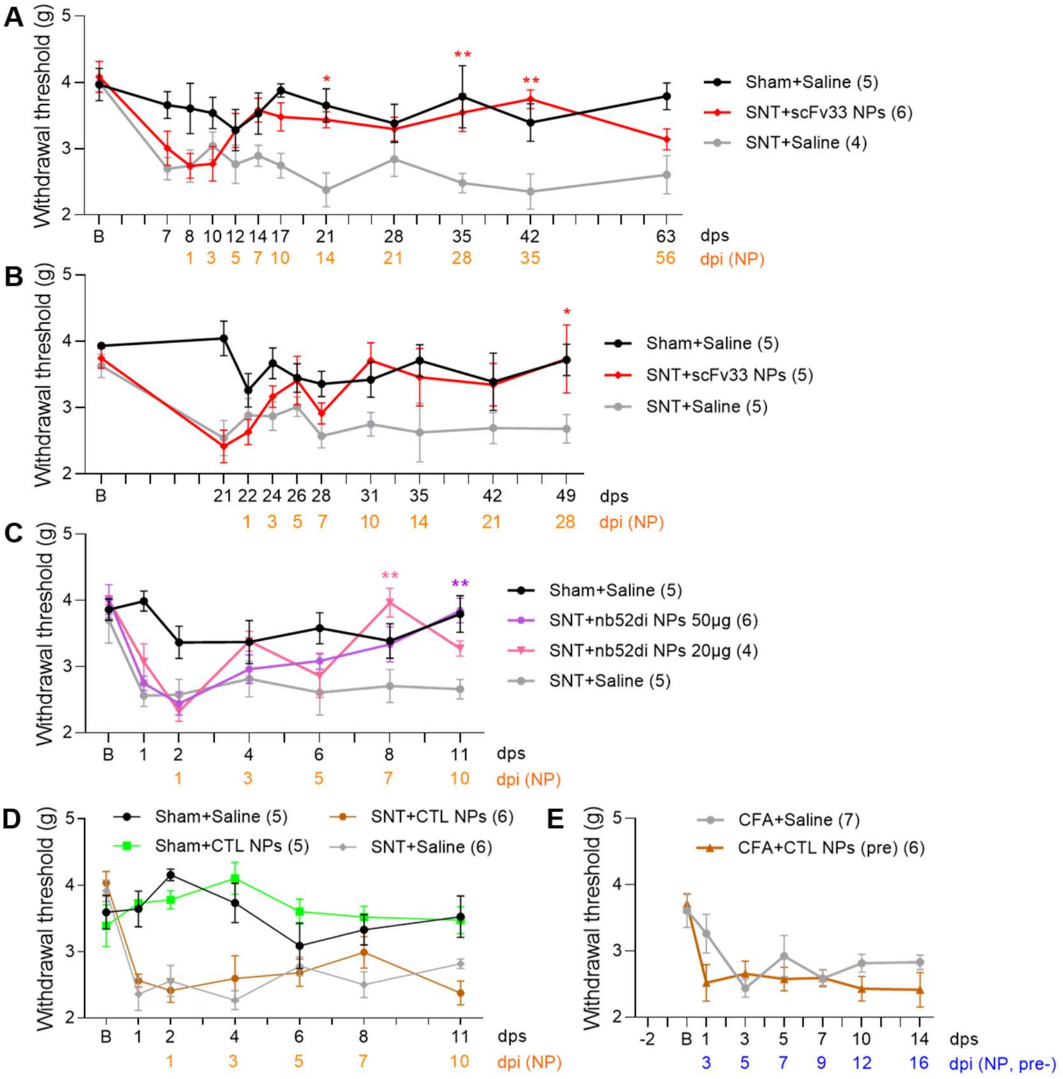


**Supplementary Figure 8. Extended validation of the analgesic effects of PLGA nanoparticle–mediated anti-TLR2 antibody gene delivery across doses and timepoints. (A,B)** Mechanical withdrawal thresholds in mice subjected to SNT surgery **(A)** or CFA injection **(B)** were measured using the electronic von Frey test following control (CTL) NP administration. **(C,D)** Mechanical withdrawal thresholds in the SNT model were measured using the electronic von Frey test following NP administration on day 7 **(C)** or day 21 **(D)** post-surgery**. (E)** Mechanical withdrawal thresholds in the SNT model were measured using the electronic von Frey test following NP administration at doses of 50 μg or 20 μg on day 1 post-surgery. All data represent the mean ± SEM. Numbers in parentheses indicate the number of animals per group. Statistical significance was determined by two-way ANOVA with Tukey's post hoc test (*p < 0.05, **p < 0.01 vs. SNT + saline group). dps, days post-surgery; dpi, days post-injection.

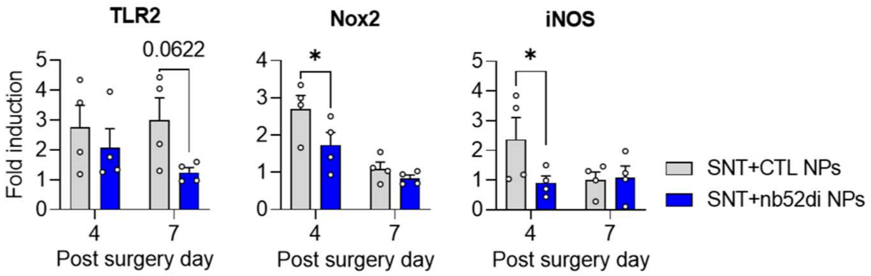


**Supplementary Figure 9. Additional proinflammatory gene expression analysis following PLGA nanoparticle–mediated anti-TLR2 nanobody gene delivery.** SNT-subjected mice were intrathecally injected with nb52 dimer (nb52di) or control (CTL) NPs (200 μg) on day 1 post-surgery. On day 4 or 7 post-surgery, lumbar spinal cord tissues (L4–L6) were harvested and analyzed by qRT-PCR to assess mRNA expression levels of additional inflammation-related genes. Expression levels of TLR2, Nox2, and iNOS are shown relative to the saline-injected sham group. Each dot represents an individual mouse. Data represent the mean ± SEM. Statistical analysis was performed using two-way ANOVA followed by Tukey's post hoc test (*$p < 0.05$ vs. CTL NP group).

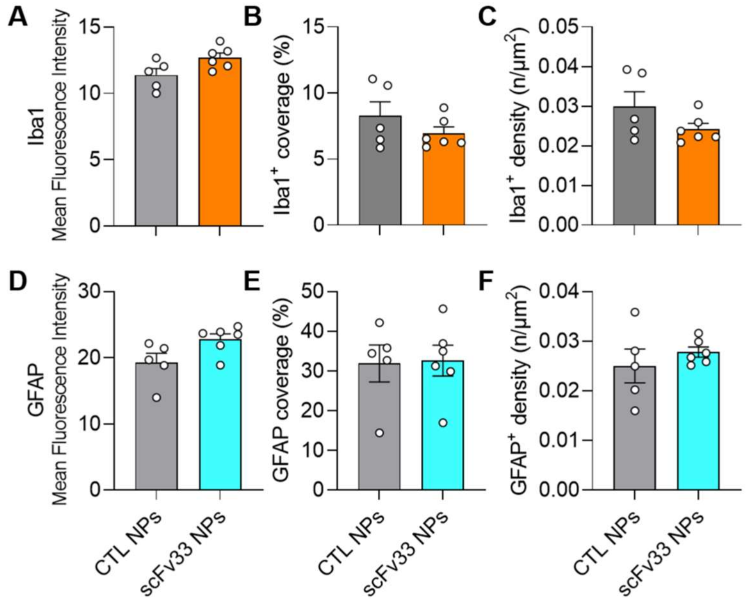


**Supplementary Figure 10. Quantitative fluorescence-based analysis of glial activation following scFv33 nanoparticle treatment.** Quantification of Iba1$^+$ microglia and GFAP$^+$ astrocytes in the spinal dorsal horn of SNT-subjected mice treated with control (CTL) or scFv33 NPs (200 μg). All analyses were performed on day 7 post-surgery. **(A–C)** Quantification of Iba1 immunofluorescence intensity **(A)**, coverage **(B)**, and cell density **(C)**. **(D–F)** Quantification of GFAP immunofluorescence intensity **(D)**, coverage **(E)**, and cell density **(F)**. Each dot represents an individual mouse. Data represent the mean ± SEM. Statistical analysis was performed using two-way ANOVA followed by Tukey's post hoc test.
